# Quantitative measurements of hydroxyl radicals generated by irradiated titanium dioxide nanoparticle suspensions

**DOI:** 10.1101/2019.12.27.889618

**Authors:** Jason A. Coral, Christopher L. Kitchens

**Affiliations:** Clemson University; Clemson, SC, USA

**Keywords:** TiO_2_ nanoparticles, phototoxic, NOM, hydroxyl radical, attenuation

## Abstract

Increased use of titanium dioxide (TiO_2_) nanoparticles in different applications has increased risk for adverse environmental implications based on an elevated likelihood of organism exposure. Anatase TiO_2_ is photoactive with exposure to ultraviolet light. TiO_2_ nanoparticle exposure to UV-A radiation in aquatic environments generates hydroxyl radical species, which may ultimately be responsible increased organism toxicity. The present research demonstrates that the rate of radical generation heavily depends on exposure conditions, particularly the presence of natural organic matter (NOM). Environmentally relevant concentrations of TiO_2_ nanoparticles were co-exposed to increasing NOM amounts (measured as concentration of dissolved organic carbon (DOC)) and UV-A intensities. Hydroxyl radical generation rate was determined using fluorescence spectroscopy. Radical generation rate was positively correlated to increases in TiO_2_ concentration and UV-A intensity, and negatively correlated to increased DOC concentration. Nanoparticle aggregation over time and decrease in light transmission from NOM had negligible contributions to the generation rate. This suggests the decreased radical generation rate is a result of radical quenching by NOM functionalities. *D. magna* toxicity to hydroxyl radicals is also demonstrated to decreased following the addition of DOC. These results correlate with the rate generation data, indicating that DOC provides rate attenuation that is protection to organisms. These conclusions demonstrate the importance considering exposure conditions during TiO_2_ toxicity testing, and during TiO_2_ waste management and regulatory decisions.

## INTRODUCTION

Anatase titanium dioxide nanoparticles (TiO_2_ NPs) are an active component in a multitude of industrial, personal, and everyday products. Specific uses include photocatalysts, paints, and surface coatings[1] due to the intrinsic abilities to generate excited electrons when exposed to ultraviolet-A (UV-A) radiation. The photo-induced mechanism of electron promotion generates free radicals, which has found application in the generation of H_2_ gas, decomposition of organic molecules and persistent chemicals, and killing of bacteria and other microorganisms[2,3].

Growth in use of TiO_2_ NPs is such that Robichaud et al. estimates that upper bound production of TiO_2_ NPs will surpass production of bulk TiO_2_ by 2022. By 2025, worldwide production of TiO_2_ NPs is expected to exceed 2 million metric tons per year [4]. Lazareva et al. (2014), predicts that approximately 88,000 tons of TiO_2_ NPs per year will be released (either through natural processes, disposal, or accident) to global environmental compartments[5]. Of this, almost 50,000 metric tons are estimated to escape wastewater treatment processes (WWTP) in the effluent, although the efficacy of removal is often dependent on the WWTP global location. WWTP with modern treatment methods are expected to sequester close to 97% of TiO_2_ NPs in biosolids. However, many WWTP use antiquated technology and the biosolids are often reutilized as fertilizer, potentially reintroducing the nanoparticles into freshwater systems[5]. Additional TiO_2_ NP loads to water systems from surface coating/paint runoff are most likely more problematic in urban areas. As predictions of exponential increases in NP manufacturing over the coming years, increasing amounts of nanoparticles can be expected to reach environmental compartments.

The movement of NPs to environmental compartments demonstrates the hazard of TiO_2_ NPs to aquatic organisms. The risk presented by TiO_2_ NPs can be understood when considering the photocatalytic effects of the nanoparticle. UV-A radiation at wavelengths of 382 nm has the equivalent energy of 3.2 eV; the approximate bandgap energy of anatase TiO_2_ NPs. Energy of this wavelength excites electrons in the valance band, promoting them to the conductance band, and leaving behind areas of positive charges referred to as ‘holes’. Electrons are transferred from water molecules to fill the holes, generating hydroxyl radicals (OH•), and initiating oxidative reactions. The promoted electron will induce a reductive pathway reaction via transfer to oxygen molecules, generating superoxide anions (•O_2-_)[6]. This photocatalyzed radical generation degrades chemicals and kills microbes, making TiO_2_ nanoparticles attractive for biocides and surface treatments.

Fujishima (2008) details the following hydroxyl generation schemes from the oxidation of superoxide, resulting in both surface adsorbed, and free hydroxyl radicals:

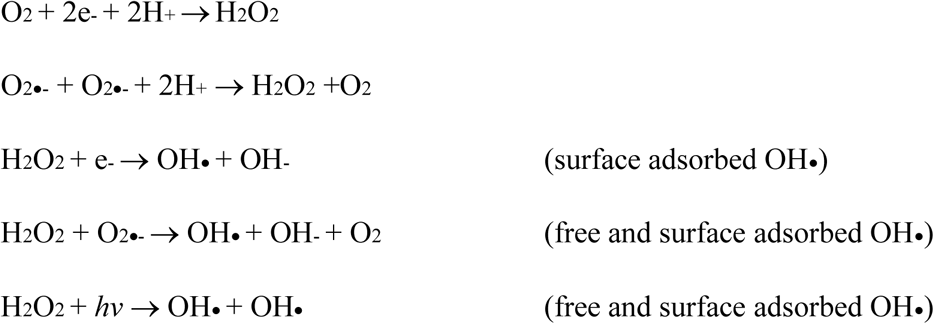

Reactive oxygen species (ROS) are well known to cause a number of biological dysfunctions through oxidative damages. Hydroxyl radicals exhibit the highest reduction potential of all ROS. These radicals are responsible for initiating lipid peroxidation cycles resulting in membrane disruption, can generate DNA adducts causing sequence mutations, and will initiate radical cycling cascades within cells [7] Superoxide anions are likewise responsible for initiating cellular response cascades, generating hydroxyl radicals in the process[7]. Taking this into consideration, it should come as no surprise that TiO_2_ nanoparticle toxicity drastically increases under UV irradiation[8].

Multiple researchers have investigated the toxicity of TiO_2_ under UV irradiation[9-13]. Ma et al.(2012), conducted a robust battery of tests showing that the phototoxicity of TiO_2_ nanoparticles to *D. magna* is partially dependent on the intensity and wavelength of UV light[14]. Li et al.(2015), demonstrated the effect of UV exposure time and intensity on TiO_2_ toxicity to *Hyalella azteca*[15]. Multiple studies indicate that the toxicity can vary by multiple orders of magnitude depending on a number of physical particle factors, such as crystal configuration[16], size of nanoparticle[17], aggregation[18], and surface coatings[16,19]. TiO_2_ nanoparticle toxicity has been shown to follow the Bunsen-Roscoe Law of Reciprocity, which describes the generation of photochemical products as being proportional to the product of light intensity and time[20]. This relationship is well demonstrated by PAHs, wherein low concentrations of chemicals and high doses of UV light produce equivalent impact as high concentrations of chemicals and low doses of UV light[21,22].

An important factor to consider is the effect of dissolved organic carbon (DOC) existing as natural organic matter (NOM). NOM is a significant component of natural water systems, comprised of humic acids, fulvic acids, lignin, proteins, and many other organic compounds. It is ubiquitous in natural waters but the particular molecular configuration and relative concentrations vary spatially and temporally[23]. Due to the persistence of NOM in water systems, it is important to determine its effects on radical generation from irradiated TiO_2_.

Until recently, literature regarding TiO_2_ nanoparticles did not focus on the more complex interactions that nanoparticles undergo in surface freshwaters and the resulting effect on toxicity. Studies have focused mainly on the effect of UV light and physical characteristics of the nanoparticle, such as size, crystalline configurations and coatings[13]. More recent work has demonstrated the necessity of incorporating NOM into studies, due to both the ubiquitous presence of NOM in all surface waters, and the inherent ability of the macromolecule to quench radicals[8,24]. Identifying the influences of changing environmental conditions on the TiO_2_ nanoparticle would help to develop a better understanding of the link between ROS generation and thus TiO_2_ toxicity. By bounding experiments at reasonable environmental concentrations of TiO_2_ and NOM, and light intensities, better predictions about the behavior of TiO_2_ NPs outside of a laboratory setting can be made. Based on previous literature, we hypothesize that variations in TiO_2_ concentration, light intensity, and NOM can be correlated with hydroxyl radical generation, which invariably influences environmental toxicity.

## MATERIALS AND METHODS

### Full Factorial Design

A full factorial approach to exposures was used for this experiment. The TiO_2_ NP concentrations tested were 0.0, 0.50, 1.00, 3.50, 5.00, 7.00, 10.5, and 14.0 mg/L. Dissolved organic carbon (DOC) concentrations were 0.00, 1.57, 2.95, 4.28, and 5.71 mg/L. UV intensities were measured as irradiant intensity per nm, averaged across 320-400 nm, and were 0.00, 2.671, 4.301, and 5.167 µW/cm^2^/nm.

### Titanium Dioxide Nanoparticle Suspensions

Anatase titanium dioxide nanoparticles (Aldrich, <25-nm, 99.7% metals basis) were suspended in 18 mega-Ohm water, at a concentration no greater than 100 mg/L. Upon the initial dispersion into stock suspensions, TiO_2_ NPs were stirred for 10 minutes and sonicated for 2 hours (in 15 minutes on/5 minute off) intervals using an immersion-tip sonicator. Before use, stock suspension was sonicated for 15 minutes, and lightly stirred during dilutions to ensure complete suspension. All suspensions were diluted with EPA recipe Moderately Hard Water (MHW) (96.0 mg/L NaSO_4_, 60 mg/L CaSO_4_-H_2_O, 60 mg/L MgSO_2_, 4.0 mg/L KCl; pH: 7.9-8.3, Hardness: 80-100, Alkalinity: 57-64) [25]. Nanoparticle size distributions were measured using Hitachi H7600 TEM. Intensity weighted hydrodynamic diameter was determined by DLS using a Wyatt Dawn Heleos-II Dynamic, at ambient temperature. UV transmission was determined, using a Varian Cary 50 Bio UV-Vis spectrophotometer, analyzed in dual beam mode from 300 to 700 nm at a scan rate of 100 nm/min, with 1-nm intervals. Aliquots of 0.5 mL were analyzed using quartz 1-cm x 1-cm x 4.5-cm plastic cuvettes (Spectrasil), at room temperature. Zeta potential was measured using a Malvern Zetasizer ZS, at room temperature. DLS and UV transmission were run for the full factorial of TiO_2_, DOC concentrations and UV intensities, measured at 0, 24 and 48 hours. Zeta potential was only determined at initial time of preparation, for the full factorial of DOC and TiO_2_.

### Natural Organic Matter

Natural organic matter was obtained directly from the Suwannee River headwaters, at Suwannee River Visitors Center in Fargo, Georgia. Water was filtered through 0.45-micron filters to remove all non-dissolved components. Filtered SWR was lyophilized to determine total amount of dissolved organic matter (DOM) per liter. Total organic carbon concentration was determined using a Shimadzu TOC-V Carbon Analyzer. Stock concentrations were directly diluted in EPA recipe MHW to achieve working concentrations.

### Light System

UV irradiance was generated using CXL Topaz 40W Blacklight Blue T-12 Fluorescent lights. Standard Lab lighting was generated using Sylvania 40W Cool White T-12 Fluorescent lights. Lights were installed in a plywood light box, measuring 48” x 12” x 12.5”, with an 8” distance from bulb to bench surface. Intensity and spectral output was measured using an OceanOptics JAZ Photospectrometer equipped with a cosine corrector. Spectroscopic data was analyzed using OceanView 1.5.2.

### Electron Paramagnetic Resonance Spectroscopy

EPR spectroscopy was used to characterize radicals produced from the irradiation of TiO_2_ NP suspension. EPR measurements at the X-band were acquired using a Bruker EMX spectrometer, with a quartz flat cell inserted directly into the microwave cavity at ambient temperature. For all experiments, the following parameters apply: Modulation frequency and amplitude were 100 kHz and 0.5 G, microwave frequency was 9.759 GHz, microwave power was 1.00 mW. Time constant and conversion time equaled 81.92. Sweep width was 100 G centered at 3479 G. The g-factor of 2,2,-diphenyl-1-pierylhydrazyl (DPPH; g=2.0036) was used as reference. Approximately 60-90 seconds elapsed between sample addition to flat cell and insertion. Spectra were analyzed using WIN EPR software.

5,5-Dimethyl-1-pyrroline-N-oxide (DMPO) was used as a spin trap for both standards and samples. Copper sulfate/ascorbic acid/hydrogen peroxide standards were used to verify the EPR hydroxyl radical signature. CuSO_4_ (15 uL, 300 µM), ascorbic acid (9.38 uL, 375 µM), 3-morpholinopropane-sulfonic acid (MOPS) buffer (50 uL, 10 µM), H_2_O_2_ (11.25 uL, 22.5 µM), and ultra-pure deionized water (414.38 uL) were combined in a microcentrifuge tube and mixed. DMPO (25 uL, 25 mM) was immediately added, mixed, and the solution was transferred to the quartz flat cell and immediately inserted into the EPR microwave cavity.

TiO_2_ suspensions were made in 50-mL volumetric flasks and separated into triplicate scintillation vials of 15-mL per vial and covered with UV transparent Aclar® film. Samples were irradiated under CLX black lights for 48 hours, with analysis points at 0, 24, and 48-hr. At each time point, the DMPO was added to the samples, and the vials were removed from the UV box. Aliquots of 0.5 mL for each sample were taken, added to the quartz flat-cell and immediately inserted into the microwave cavity for analysis.

### Fluorescence Spectroscopy

Fluorescein dye was prepared by dissolving 0.047 g HEPES buffer (Alfa Aesar, 99%) into 10 ml ultrapure DI water in a scintillation vial. 500 uL of 10M KOH (Acros, 85%) was added and stirred. 0.0367 g of fluorescein, free acid (Sigma) was added and stirred until complete dissolution occurred. The solution was acidified with 130 µL HNO_3_ (BDH, 69%) and adjusted to pH 7.5. Final stock concentration was 100 mM. Working concentration did not exceed 7.5 µM. Sample dye concentrations were varied depending on the range of hydroxyl radical generation.

Fluorescence analysis was performed on a Horiba Fluoromax-4 fluorescence spectrometer. Excitation wavelength was 467 nm and emission intensity was measured from 400 to 700 nm. Excitation slit width was 1.0 nm; emission slit width was dependent on initial concentration of fluorophore.

Calibration curves were generated using horseradish peroxidase and hydrogen peroxide, as per the Hydroxyl Radical Antioxidant Capacity (HORAC) assay[26]. 5 mg of horseradish peroxidase (HRP) (Sigma, lyophilized powder, AU: 310 units/L) was dissolved in 5 mL of 18 mega-Ohm DI water to give solution of 310 AU/L. Hydrogen peroxide (BDH, 30%) concentrations (0 to 10 µM) were combined with 107.6-uL of HRP solution, 150 uL of 1M potassium phosphate buffer (BDH, 98%) and requisite amount of fluorescein. Solutions were then diluted to 3 mL. Separate calibration curves were made for each dye concentration, with new calibration curves made for each new stock solution. To determine concentration (µM) of hydroxyl radicals at a given emission, the concentration of H_2_O_2_ is multiplied by two, as horseradish peroxidase generates hydroxyl radicals in stoichiometric proportions. Calibration curves are shown in **Supplemental Information.**

All sample solutions were made in 100 ml volumetric flasks. TiO_2_ was diluted to testing volume (0.00, 0.500, 1.0, 3.5, 5.0, 7.0, 10.5. or 14 mg/L) along with DOC (0.00, 1.57, 2.95, 4.28, 5.71 mg/L) and fluorescein dye (1.5, 5.0, 7.5, 10.0 µM/L); 30 ml was dispensed into beakers, in triplicate. A 3 ml aliquot was removed from each for analysis, and these suspensions were immediately covered with Aclar ® Fluoropolymer film, a UV transparent film, and placed in the lightbox. Dark control 0 µW/cm^2^/nm samples were additionally covered with aluminum foil to ensure the complete blockage of UV irradiation. The samples were only removed to take 3 mL for analysis every 12 hours.

Hydroxyl radical generation was determined by measuring the difference in emission from time zero. All concentrations were run in triplicate, and each sample scan was an averaged triplicate scan. Rates were determined from the slope of a linear regression line for each concentration across 48 hours. 0 mg/L TiO_2_ concentrations were treated as a blank and generation rates were subtracted to account for decreases in emission not related to hydroxyl radical generation.

### Statistical Analysis

All statistical analysis was performed using JMP software (version 12.1.0). Residuals confirmed for normativity using the Sharpiro-Wilk test. Comparative analysis of z-average aggregate group means was performed using analysis of variance (ANOVA), with Tukey-Wagner HSD as a *post hoc* for all treatments demonstrating statistically significant differences between levels. Comparative analysis was performed on all rate data using a comparative analysis of means with a Sidak adjustment, and Tukey’s HSD to determine significant differences between rates (α=0.05).

## RESULTS and DISCUSSION

### Natural Organic Matter Characterization

Triplicate measurement determined a concentration of 107.46 (±19.84) mg/L DOM in solution. Dissolved organic carbon was determined to be 61.34 (±0.43) mg/L by Total Organic Carbon analysis. HPLC-ICP determined metal content is shown in **Supplemental Information.**

### Lighting Characterization

Light intensities generated by CLX black lights in the UV-A range (320-400 nm) measured between 2.671 μW/cm2/nm to 5.188 μW/cm2/nm, with peak intensity occurring at 365 nm. These light intensities can be compared to the intensity of sunlight at noon in South Carolina, 45.08 μW/cm2/nm. Lighting characterization can be found in the Supplemental Information section. The UV-Vis percent transmission spectra were examined to measure light impedance as a result of increasing DOC. Within the working range of the experiment (0 mg/L DOC to 5.71 mg/L DOC), the increases in DOC concentration result in a maximum of 10% transmission decreases. Concentrations of DOC at 16 mg/L to 60 mg/L result in 16-60% UV attenuation. This data is summarized in the Supplemental Information. These results agree with Wormington et al. (2017).

### TiO_2_ Suspension Characterization

The nanoparticles used in this experiment were industrial grade nanoparticles with a wide distribution range. Primary particle size was measured by TEM to be 21 ± 19 nm. The greatest distribution density of nanoparticles was 14 nm. A histogram can be found in the **Supplemental Information.** Although these nanoparticles were difficult to measure using conventional TEM techniques due to aggregation, size conforms to product specifications.

The initial zeta potential was measured at 0 hour using a Malvern Zetasizer. No trends were evident across the TiO_2_ gradient at any DOC concentration, nor across the DOC gradient at any one TiO_2_ concentration. Zeta potential ranged from ∼11 mV to ∼20 mV. These data is found in the Supplemental Information.

The z-average hydrodynamic diameter of the aggregates was measured using dynamic light scattering. The z-average hydrodynamic diameter of TiO_2_-aggregate size varied as TiO_2_ and DOC concentrations were increased, with a general trend of higher TiO_2_ concentrations resulting in larger aggregations. 0-hour aggregate measurements ranged from 60 nm to 470 nm, 24-hour measurements ranged from 28 nm to 623 nm, and 48-hour size measurements ranged from 5 nm to 454 nm. Increases in light intensity resulted in decreases in z-average diameter particularly at the highest light intensities (5.177 μW/cm2/nm), in the lowest concentrations of TiO_2_. One-way Analysis of Variance (ANOVA) was run on all categories (TiO_2_, DOC, UV_i_) to determine if there were any statistically significant differences in group mean aggregate sizes. Any ANVOA found to show statistically significant difference was followed by a Tukey-Kramer HSD *post-hoc* analysis to determine which categories showed statistically significant group mean z-average diameter differences. This data can be found in the **Supplemental Information.**

The variation of these aggregate sizes can be attributed to the multiplicity of interactions occurring between the TiO_2_ NPs, natural organic matter, and irradiation as time passes. Increases of TiO_2_ concentration will result in larger aggerates, a result of charged surface interactions of the nanoparticle[27,28]. These larger aggregates experience gravimetric settling over time, resulting in removal from the water column[9]. However, NOM coated nanoparticles will experience changes in surface charge, further affecting how these particles interact. NOM has an overall negative charge and has been shown to stabilize nanoparticles in solution[24], possibly resulting in larger aggregates staying in suspension.

The steric interactions of larger NOM molecules will contribute to colloidal stability, resulting in the formation of smaller aggregates at higher concentrations. Loosli et al. (2013), reported nanoparticle aggregation enhanced by humic acid (a component of NOM), with maximum aggregate size occurring at 2.0 - 3.0 mg/L humic acid, with rapid size decrease at concentrations above 3.0 mg/L[27]. However, this was not examined in the presence of ultraviolet irradiation. Advanced oxidation of NOM functionalities, such as that produced by irradiated TiO_2_ will also modify the surface charge and steric interactions[29]. Surface charge was not measured past 0 hours due to instrument and time constraints, so we cannot conclusively state that changes in surface charge are responsible for these aggregate size changes. Inasmuch, determining the particular effect of environmental parameter changes on changes in z-average aggregate size is outside the scope of the presented research.

Overall, there is a high degree of polydispersity, contributed from NOM molecules of varying identities and sizes, nanoparticle aggregates, and buffer salts. The complexity of these suspensions demonstrate the multiplicity of interactions TiO_2_ nanoparticles undergo in an aquatic environment and demonstrate the need for further investigation.

### Identification of Hydroxyl Radical

EPR spectra indicated the formation of hydroxyl radical, superoxide anions and free electrons. A standard spectra generated with copper sulfate/ascorbic acid/hydrogen peroxide **(Figure 1A)** was compared to spectra from the irradiated sample **(Figure 1B)**. The DMPO-OH• adduct generates a clear 1:2:2:1 quartet, plainly recognizable in the standard. Spectra from the irradiated TiO_2_ suspensions demonstrates hydroxyl radical generation (confirmed by the standard CuSO_4_/Citric acid/H_2_O_2_ spectra), as well as that of superoxide radical (confirmed by literature spectra[30]). The spectra were comparable to published irradiated TiO_2_ spectra by a number of researchers[31-33]. Although both superoxide and hydroxyl radical were detected in the suspensions, the generation of ROS can be described entirely as hydroxyl radical for a number of reasons. These include concentration of radicals, empirical chemistry concerning hydroxyl radicals produced by irradiated TiO_2_ suspensions, and the importance of the biological effects of hydroxyl radicals.

**Figure 1:**
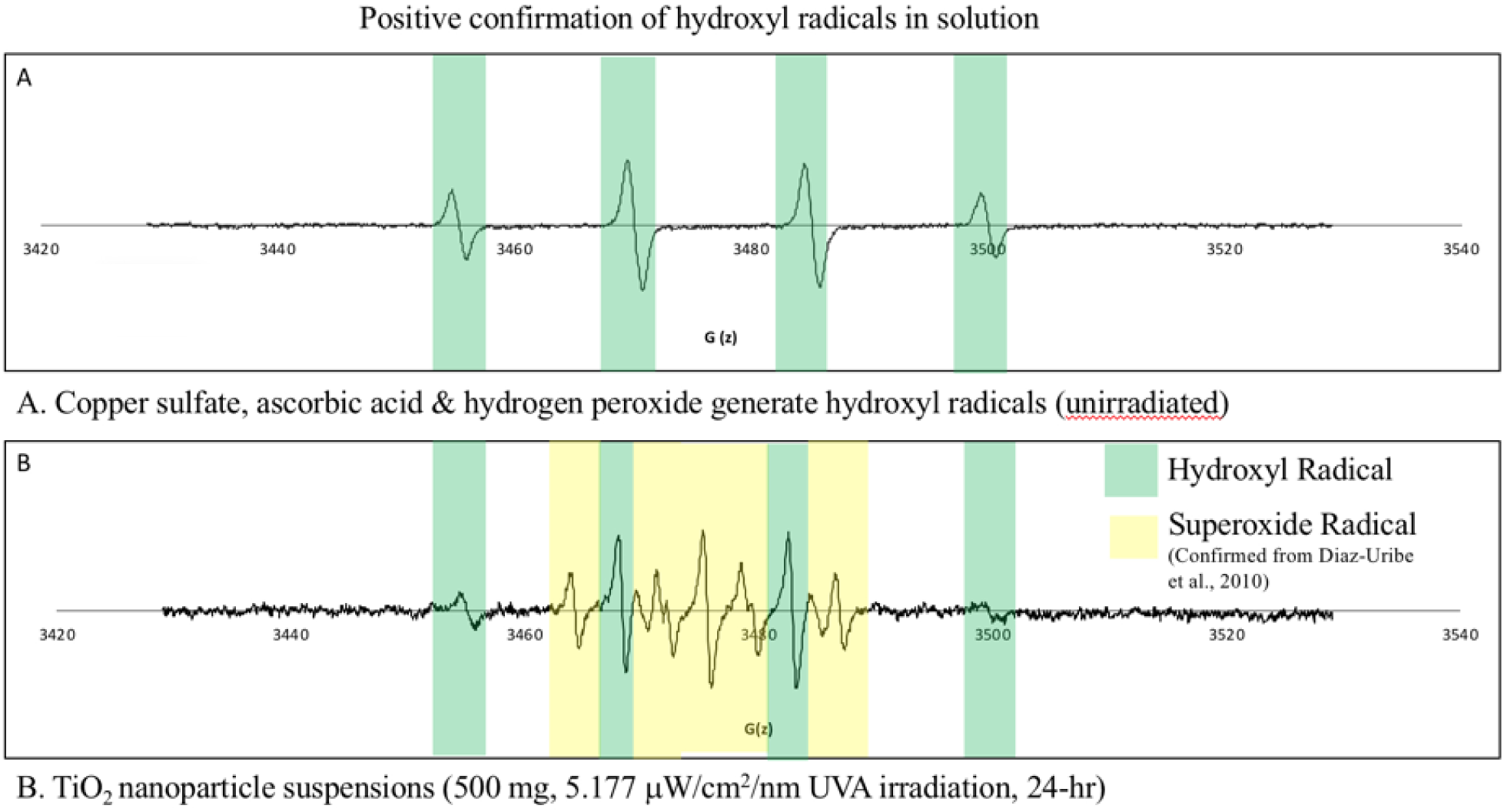
Spectra of ROS Characterized by Electron Paramagnetic Resonance Spectroscopy. EPR characterization of hydroxyl radical formation. Characterization of radicals formed in irradiated nano-TiO_2_ suspensions. A) copper sulfate, ascorbic acid and hydrogen peroxide generate hydroxyl radical without irradiation. B) TiO_2_ nanoparticles under ultraviolet irradiation generate a hydroxyl radical signature, along with signatures for superoxide. DMPO was used as a spin trap to generate both EPR spectra.

TiO_2_ nanoparticle irradiation produces majority superoxide (O_2•-_) and hydroxyl radicals (OH•). Although the superoxide radicals produced by the photocatalytic reaction have higher quantum yields than hydroxyl radicals[34,35], multiple researchers show that superoxide more readily acts as an oxidizing agent for H_2_O, and catalyze the formation of H_2_O_2_. H_2_O_2_ will then readily react with free electrons to generate OH•, with a cycling reaction possible, generating more hydroxyl radicals [6,32,35-39].

Further evidence for describing radical generation solely as the hydroxyl radical is provided by observing organic intermediates in reactions photocatalyzed by TiO_2_ and comparing these intermediates to those produced by reactions with known hydroxyl radical sources, which are consistent[40,41]. These radical intermediates include DMPO-OH adducts, of which there is significant evidence indicating majority hydroxyl radical generation [31,32,39,40,42]

Observations of photocatalytic reactions run in water-free, aerated organic solvent (D_2_O) show significant decrease in oxidation rates of organic molecules compared to reactions run in water[40]. This was further characterized by (CH_3_)_2_CHOH lack of complete degradation to CO_2_, which occurs in water/TiO_2_ reactions. Replacing H with D in the organic molecule, while retaining the aqueous solvent showed no rate reduction. This implies that the reaction is limited by the formation of active oxygen species. O-D bonds have lower ground state energies, that are not able to be overcome[40].

Finally, toxicity driven by TiO_2_ photocatalysis generally indicates the importance of ROS species. Both radical species produced (superoxide, hydroxyl radicals) are capable of inducing cellular antioxidant effects, however, superoxide is biologically unstable and generally has poor biological reactivity[7,43]. Intracellular superoxide will be readily catalyzed by superoxide dismutase (SOD) to hydrogen peroxide, which is then reduced to water and oxygen radicals via catalase or glutathione peroxidase[7]. Regardless, the hydroxyl radical is much more biologically disruptive, reacting with every type of biomolecule and nucleic acids[7,43,44]. There are no known enzymatic reactions that can scavenge OH•. Instead, antioxidants are the sole source of removal without reaction with a biomolecule[7,43]. Because the TiO_2_ nanoparticle must be activated by light outside the cell, hydroxyl radical interactions with the outer cellular membrane resulting in lipid peroxidation are the most common cause of toxicity[45]. Singlet oxygen is catalyzed by reactions of superoxide with trapped holes and has shown slight influence on toxicity, demonstrated by increased production in lipids[39,45], but ^1^O_2_ short lifetime compared to the hydroxyl radical (2 μs and 10 μs, respectively) makes this possibility less reasonable[39]. Given the numerous mechanisms of hydroxyl radical generation, the wealth of data implying that hydroxyl radicals are the most relevant radical produced, and the known biological implications, generalizing radical generation as hydroxyl radical is within the bounds of this study.

### Measurement of Hydroxyl Radical Generation Rates

Photocatalytic hydroxyl radical generation rate measurements showed significant differences for TiO_2_ nanoparticle suspensions with varied DOC concentrations up to ∼6 mg/L, UV irradiation up to 5.177 μW/cm2/nm, and TiO_2_ NP concentration up to 14 mg/L. All hydroxyl radical generation by TiO_2_ over 48 hours were observed to be linear. Generation followed distinct trends, where radical concentration increased with increasing TiO_2_ concentration, and UV intensity. Addition of DOC decreased the total amount of radicals generated. Reciprocity was demonstrated where lower concentrations of TiO_2_ and high intensities of UV light generated similar hydroxyl radical concentrations compared to high TiO_2_ concentrations and low UV intensities. All samples exposed to 0 μW/cm2/nm generated zero, or extremely close to zero hydroxyl radicals. The calculated rates are expressed graphically in **Figure 2A to 2D. Statistical summary can be found in the Supplemental Information**

**Figure 2A:**
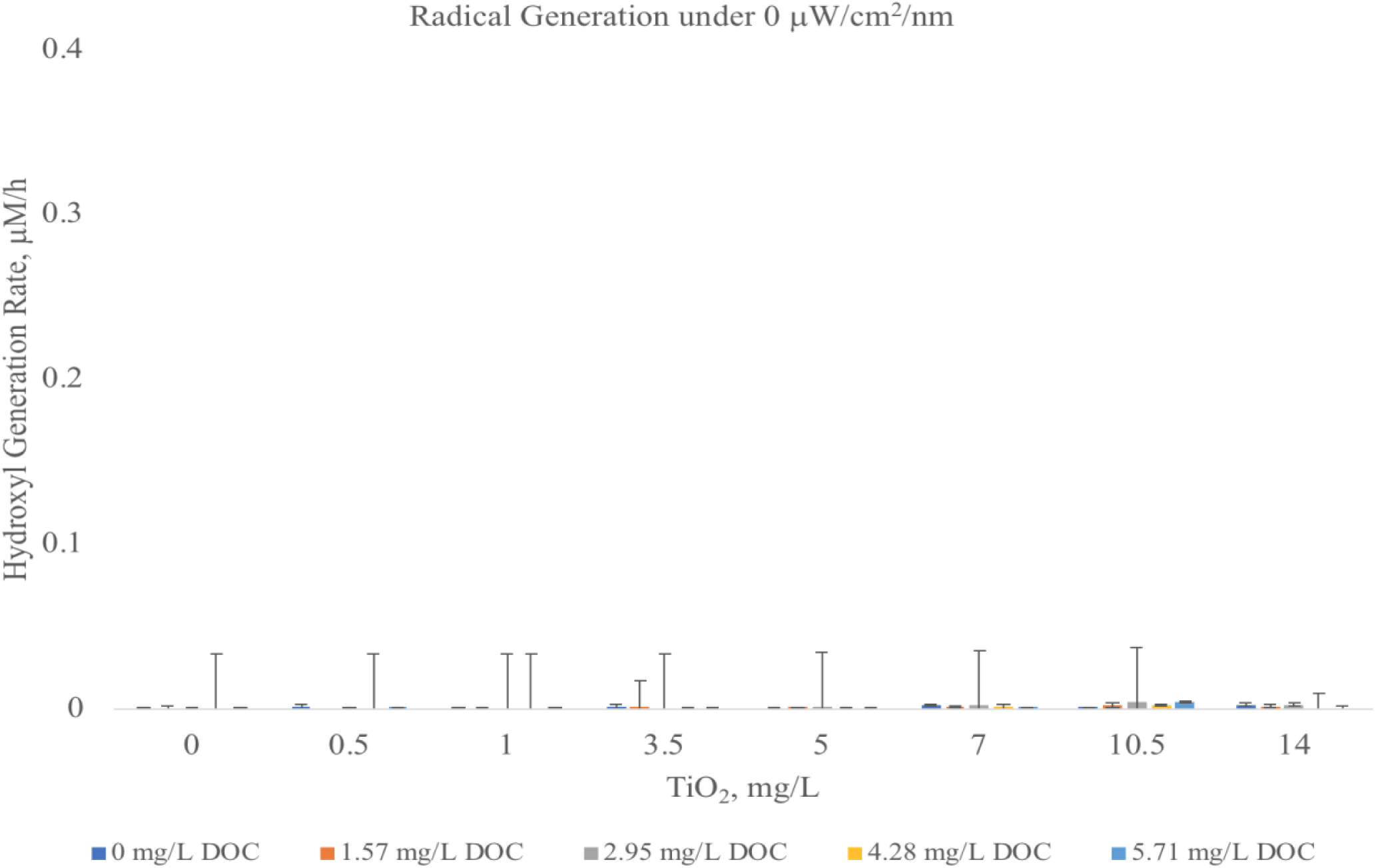
Hydroxyl Radical Generation Rates under 0 μW/cm^2^/nm. Radical generation rate as measured by fluorescence spectroscopy. Letters indicate level of significance; non-matching letters are statistically significantly different within the TiO_2_ treatment (α = 0.05, *p* < 0.05). There were no significant differences in any of the 0 μW/cm^2^/nm measurements.

**Figure 2B:**
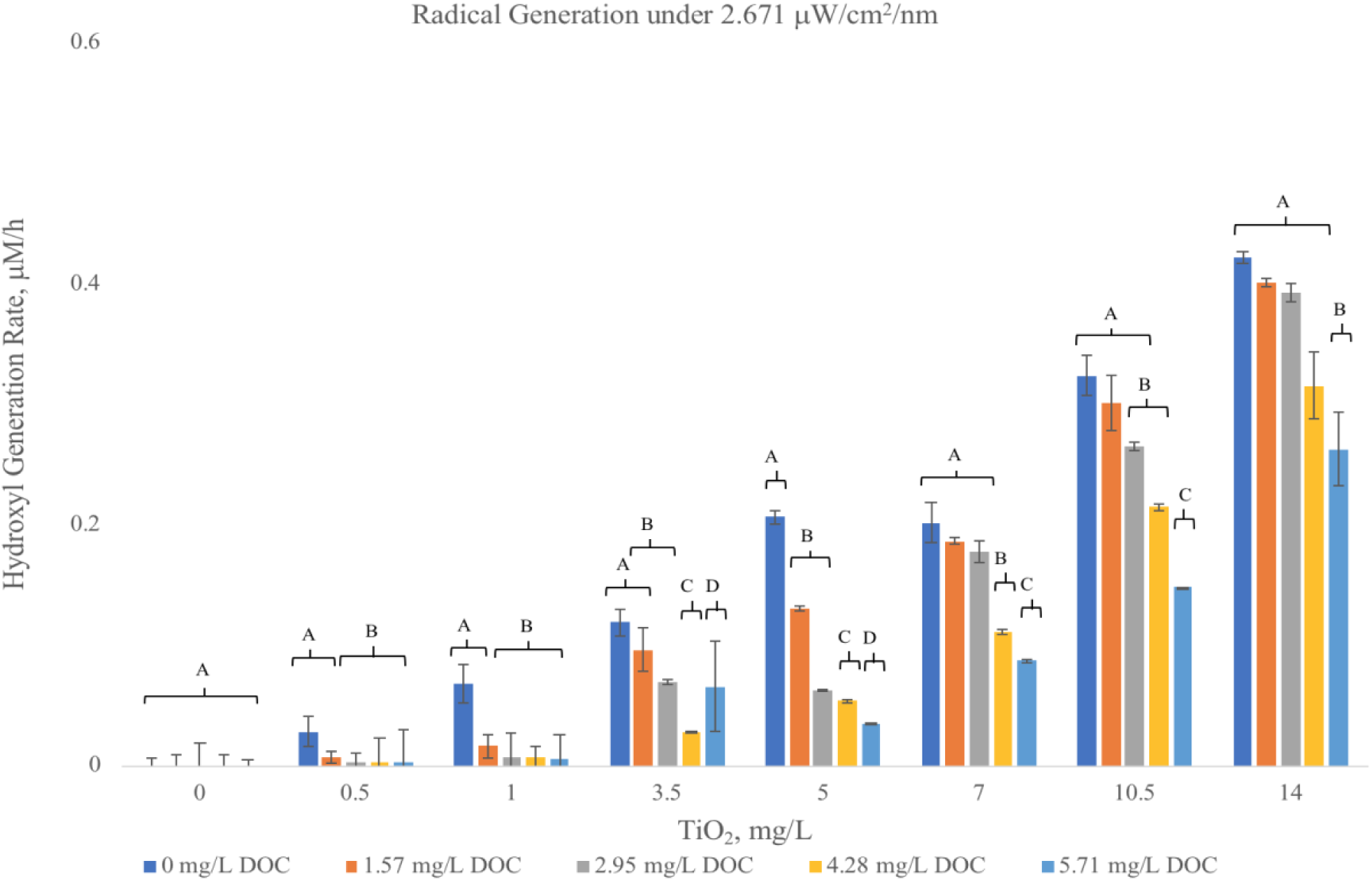
Hydroxyl Radical Generation Rates under 2.671 μW/cm^2^/nm. Radical generation rate as measured by fluorescence spectroscopy. Radical generation rate increased as TiO_2_ concentration increased and UV_i_ increased. Generation rate decreased as DOC concentrations increased. Letters indicate level of significance; non-matching letters are statistically significantly different within the TiO_2_ treatment (α = 0.05; *p* < 0.05).

**Figure 2C:**
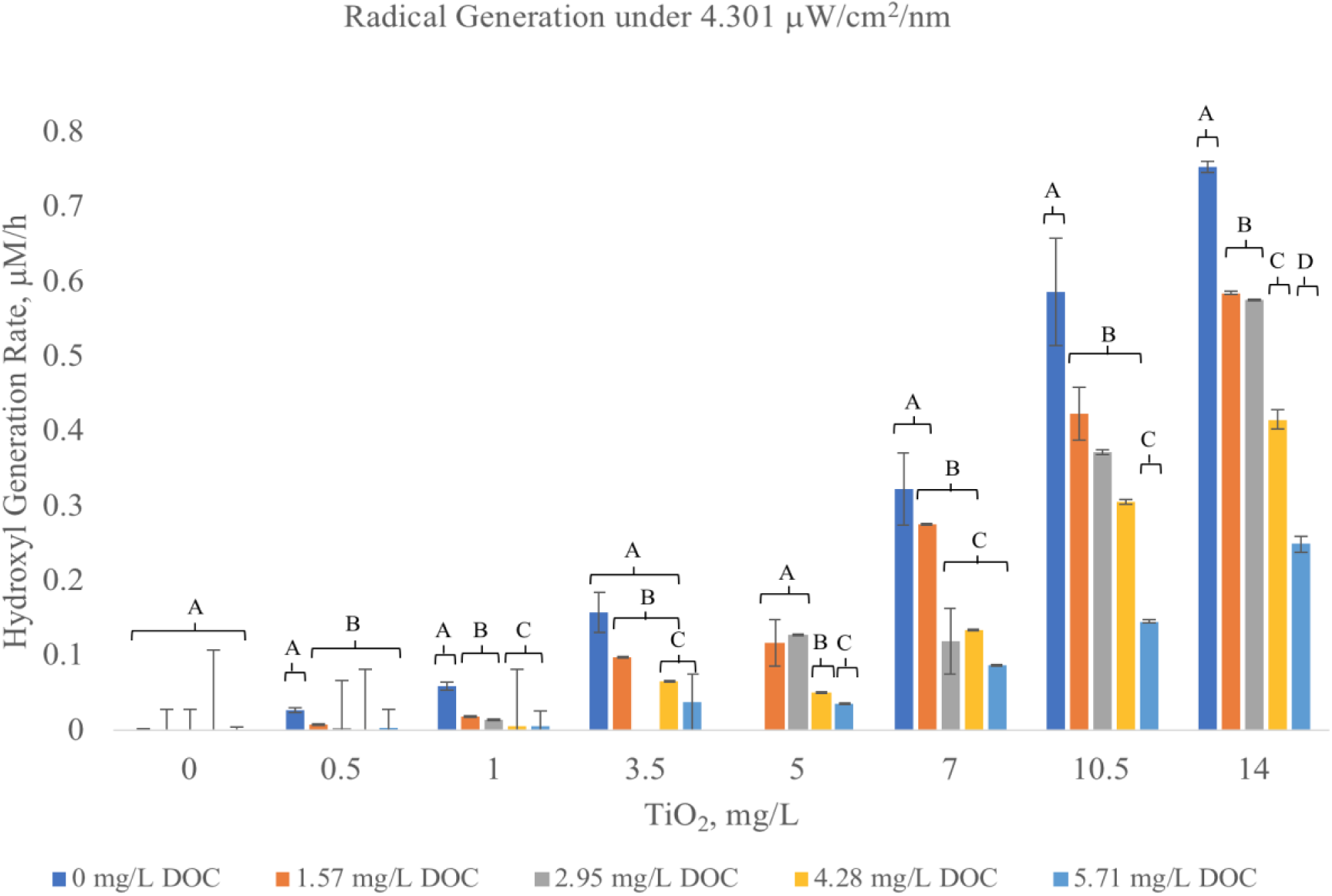
Hydroxyl Radical Generation Rates under 4.301 μW/cm^2^/nm. Radical generation rate as measured by fluorescence spectroscopy. Radical generation rate increased as TiO_2_ concentration increased and UV_i_ increased. Generation rate decreased as DOC concentrations increased. Letters indicate level of significance; non-matching letters are statistically significantly different within the TiO_2_ treatment (α = 0.05, *p* < 0.05).

**Figure 2D:**
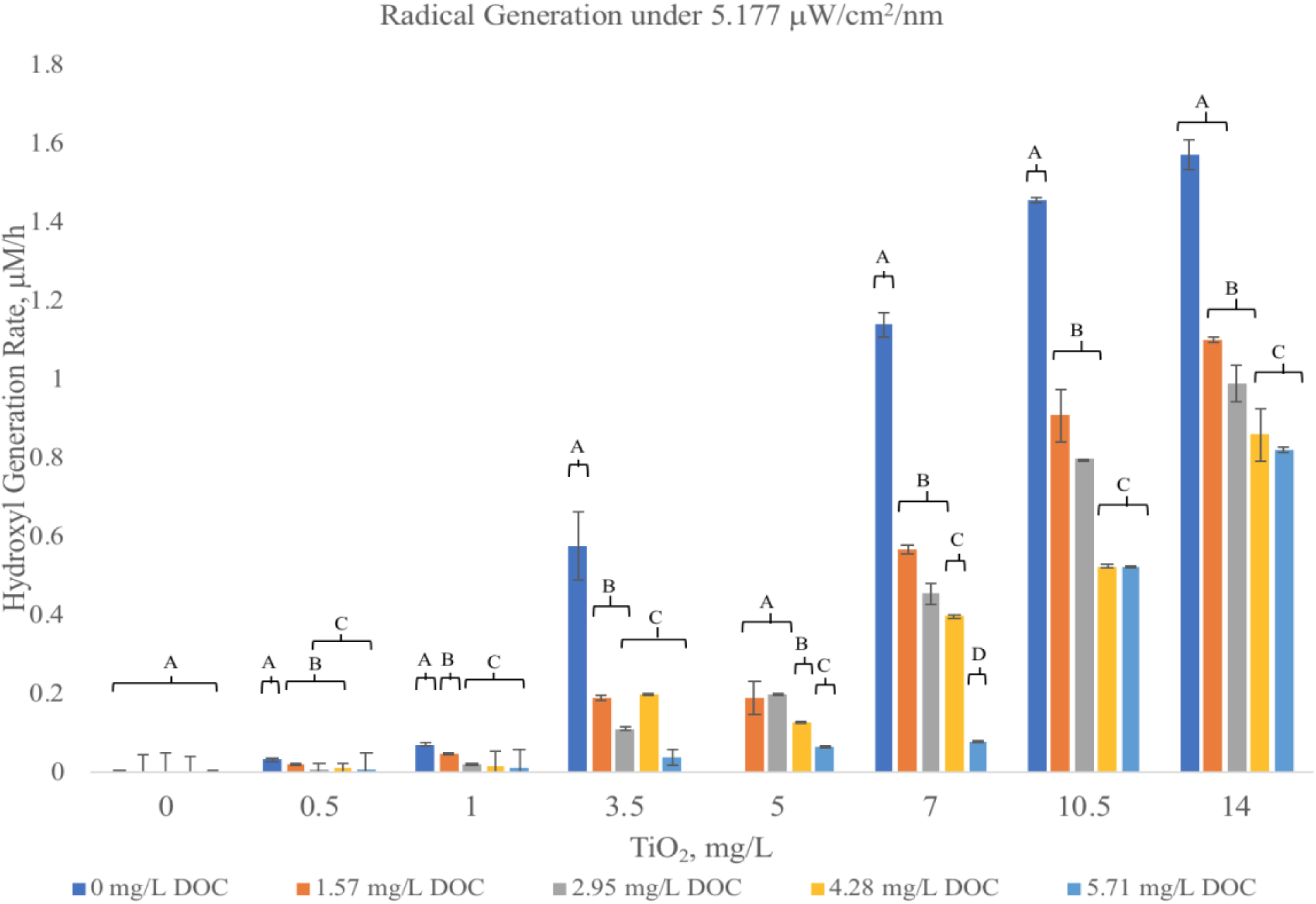
Hydroxyl Radical Generation Rates under 5.117 μW/cm^2^/nm. Radical generation rate as measured by fluorescence spectroscopy. Radical generation rate increased as TiO_2_ concentration increased and UV_i_ increased. Generation rate decreased as DOC concentrations increased. Letters indicate level of significance; non-matching letters are statistically significantly different within the TiO_2_ treatment (α = 0.05, *p* < 0.05). Statistic tables can be found in the Supplemental Information.

Radical generation was the greatest with the highest TiO_2_ concentrations (14 mg/L) with 0 mg/L DOC, under higher UV-A light intensities. These exposures consistently generated such excess hydroxyl radicals that the fluorescent dye used to measure generation was routinely diminished around the 36-hour mark, such that increased concentrations of dye were required. These results agree with expected dependence of radical rate on TiO_2_ concentration. Increased concentrations allow for increase photon/nanoparticle interaction, resulting in an increased rate of hydroxyl radicals generation. These results are in agreement with literature[6,46].

Rate generation increased as light intensity was increased. Increased intensity results in an increase in photons impinging on the surface of nanoparticles. This reaction generates more electron-hole pairs, ultimately resulting in an increase in hydroxyl radical generation, and are in agreement with literature[14,15,42,47]. Reciprocity effects were apparent in samples; exposures with low concentrations of TiO_2_ exposed to high UV intensities had similar rates of high concentrations of TiO_2_ exposed to low intensities of light. The low intensities used are comparable to those in deeper parts of the water column[48]. The significance of the radical generation at these low intensities should not be understated. The intensities studied can be expected to occur at deeper water columns depths. Increased rates of hydroxyl radical generation from TiO_2_ irradiation occurring in upper levels of the water exposed to higher UV intensities should be expected.

Rate generation significantly decreased on the addition of natural organic matter, at all concentrations and under all intensities. Although there is little published data on the effect of NOM on TiO_2_ radical generation, these results are well correlated with toxicity data, showing that increased amounts of NOM and humic acid provide a protective effect against toxicity to *D. magna*[24] and to *Chlorella* sp. [49] resulting from TiO_2_ irradiation.

Overall reduction of rate on the addition of NOM may be explained through 3 different scenarios: changes to particle surface charge resulting in changes in particle presence within the irradiated water column, reduction of light transmission due to increased NOM content, or quenching of hydroxyl radicals by NOM functionalities. TiO_2_ NPs will aggregate, and settle over time, at higher concentrations settling more quickly. Keller et al. (2010), measured settling rates of multiple TiO_2_ NP concentrations and found that concentrations less than 10 mg/L stay stable in suspension, remaining in the water column for prolonged periods. Further, aggregates in DOC that slightly larger than 300-nm and smaller remained stable[28].

Such evidence suggest that lack of suspension stability is not responsible for decreases in hydroxyl generation rate in the presented research. Forman, (2015) notes that concentrations of 10 mg/L TiO_2_ NPs have sedimentation rates that range from 10^−7^ to 10^−4^ per second, with faster sedimentation occurring in waters with higher ionic strengths, demonstrating increased rates of removal in harder waters or those with higher salinity. [50]. Brunelli et al. (2013), found low settling rates (10^−6^ to 10^−5^ per second) in freshwater at 1 mg/L and lower concentrations of TiO_2_ after 50-hrs[51]. These results indicate that substantial amounts of TiO_2_ will be suspended in freshwater water column and exposed to sunlight and may therefore demonstrated low TiO_2_ concentration rates may underestimate that found in waters exposed to direct solar radiation.

UV-A is attenuated by DOC, with a hyperbolic inflection point occurring at 1-2 mg/L DOC[48]. In waters of less than 2 mg/L of DOC, UV-A has a 1% attenuation depth (that is, the depth at which 1% of surface irradiation will penetrate) of 2 to 12 meters[39]. The present research has demonstrated that UV-A irradiation is maximally attenuated at 10% at the highest concentration of DOC. This 10% decrease in attenuation does not account for the larger decrease in hydroxyl rate generation, and therefore cannot be solely responsible for the overall hydroxyl generation rate reduction. These results agree with Wormington et al. (2017), demonstrating that the minor attenuation would not be able to cause significant decreases in generation rate[24].

This leaves only the third possibility, that quenching of hydroxyl radical by NOM functionalities is responsible for the decrease in measurable generation rate. This interaction between the generated hydroxyl radical and NOM functionalities is demonstrated by the breakdown of humic acids and other NOM molecules reported by a number of authors[52-54]. These results present an important conclusion, as they demonstrate NOM’s ability to attenuate radical production, which may allow for variations in regulatory actions regarding treatment and disposal of TiO_2_ nanoparticles. The importance of understanding the interactions of TiO_2_ photocatalysis is able to break down humic acids, removing up to 80% of DOC and 90% of UV_254_ absorbance[52-54] and has been used as a method to reduce filter biofouling[55]. Additionally, drought-induced acidification (from the reoxidation of sediment sulfur) of freshwater arboreal lakes results in DOC decreases sufficient enough to increase UV penetration depth by 3-fold. Thus, the implications of TiO_2_ interaction with NOM presented in this research has relevance due to worldwide climate change events[56].

### Daphnia magna 48-hour Acute Bioassays

These hydroxyl radical rate measurements can be used to describe the effects that changes in environmental parameters have on mortality endpoints for *D. magna.* Using EPA standard 48-hour acute bioassays. **Figure 3** shows that *Daphnia magna* mortality was well correlated with increasing TiO_2_ concentration and UV intensity. *D. magna* exposure to TiO_2_ under fluorescent lighting (0.0432 µW/cm^2^/nm UV intensity) resulted in LC_50_ toxicity of 837.0 (675.2, 1370.6) mg/L TiO_2_. Exposure to UV light at a wavelength of 320-400 nm and an intensity of 4.301 µW/cm^2^/nm resulted in nearly four orders of magnitude increase in toxicity, with an LC_50_, 0.156 (0.121, 0.199) mg/LTiO_2_. Significant attenuation occurs with addition of 4.28 mg/L DOC to suspensions exposed to UV light resulting in *D. magna* LC_50_ toxicity of 9.10 (8.37, 9.89) mg/L TiO_2_.

**Figure 3:**
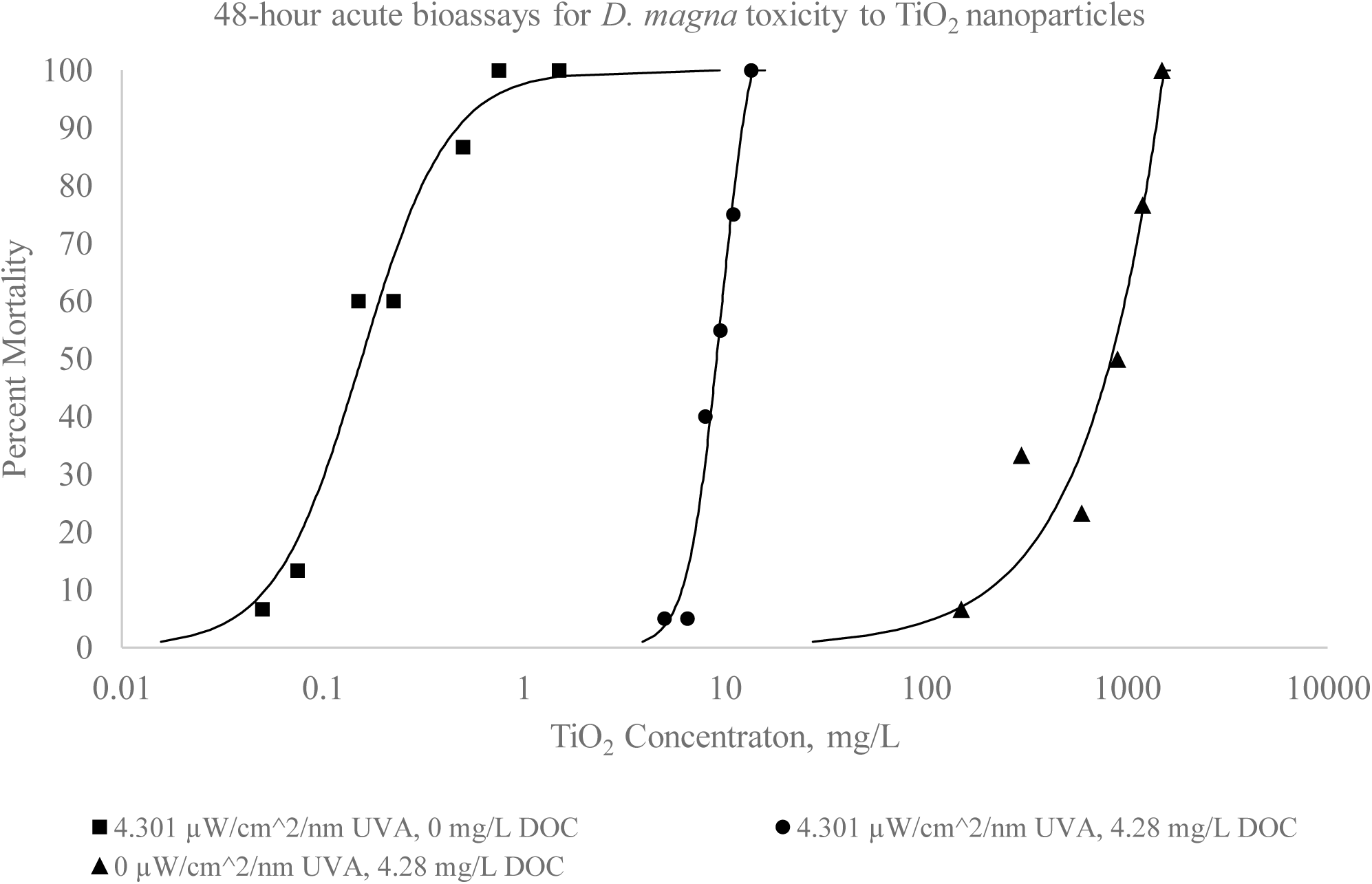
*Daphnia magna* LC_50_ toxicity to TiO_2_ nanoparticles is dependent on the presence of UV light. *D manga* 48 hour acute LC_50_ toxicity exposed to 0 µW/cm^2^/nm UV-A, 4.28 mg/L DOC: 837.0 (675.2, 1037.6) mg/L TiO_2_ NPs; 4.301 µW/cm^2^/nm UV-A, 0 mg/L DOC: 0.156 (0.121, 0.199) mg/L TiO_2_ NPs; 4.301 µW/cm^2^/nm UV-A, 4.28 mg DOC: 9.10 (8.37, 9.89) mg/L TiO_2_ NPs.

**Figure 4** shows the toxicity with UV (4.301 µW/cm^2^/nm) exposure as a function of generated hydroxyl radical. Exposures with 0 mg/L DOC demonstrated much lower hydroxyl radical LC_50_ toxicity (0.458 (0.415, 0.503) µM OH•) than in the presence of with 4.28 mg/L DOC (6.216 (5.771, 6.691) µM OH•). No hydroxyl radicals were generated under lab light conditions; thus, the data is not included in the figure.

**Figure 4:**
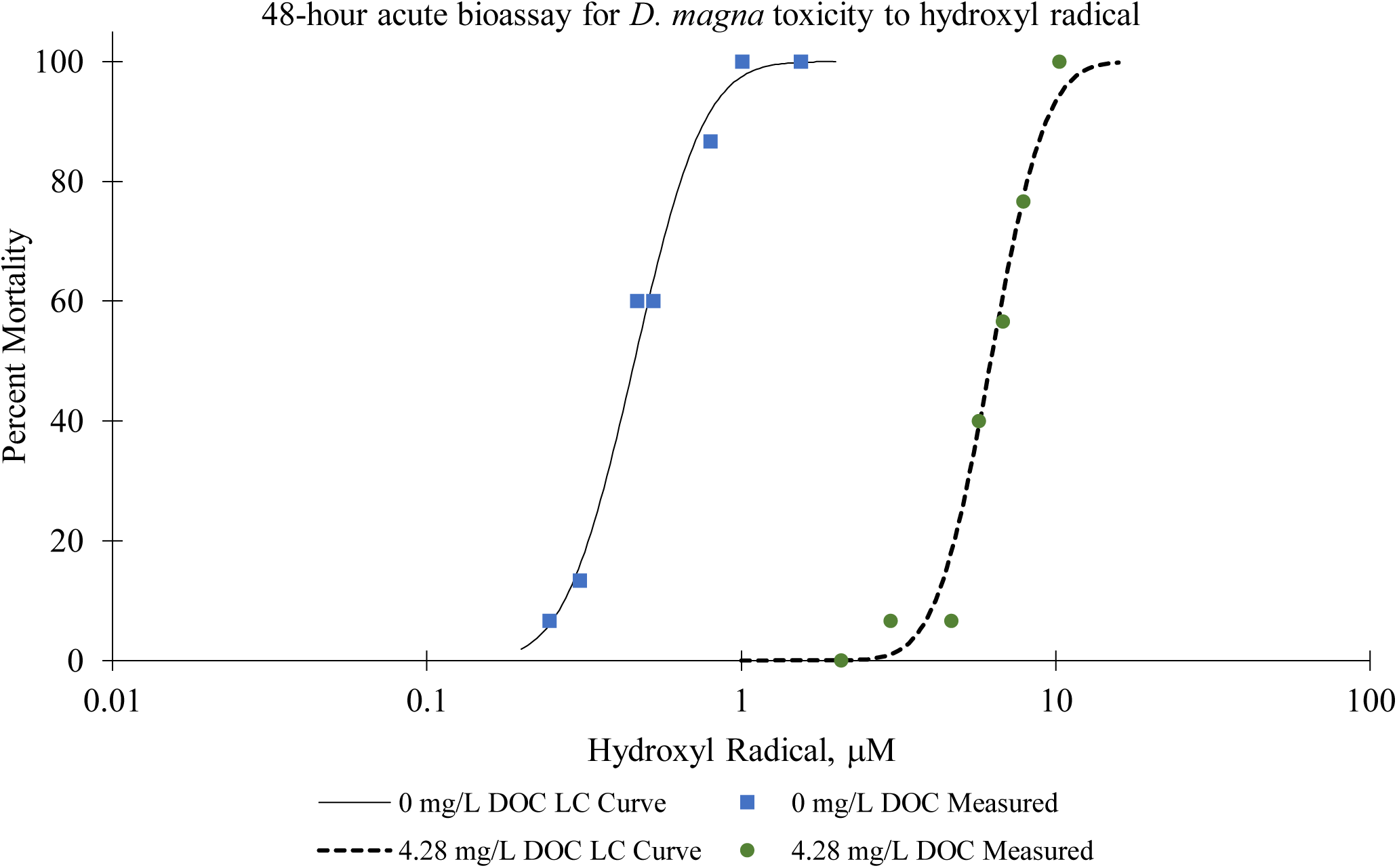
*D. magna* toxicity described in terms of hydroxyl radical. TiO_2_ generates radicals at an environmentally dependent rate. This rate is dependent on the interplay of UV intensity, DOC concentration, and TiO_2_ concentration. *D. magna* LC_50_ toxicity for hydroxyl radical at 4.301 µW/cm^2^/nm, 0 mg DOC: 0.458 (0.415, 0.503) µM OH•; 4.301 µW/cm^2^/nm, 4.28 mg DOC: 6.216 (5.771, 6.691) µM OH•.

Measuring hydroxyl radical generation under identical conditions to *D. magna* bioassays correlates toxicity to hydroxyl radicals (Figure 2.9). Conditions are also correlated to toxicity in this manner. Exposures without NOM had a rapid increase in toxicity per µM of hydroxyl radical, resulting in LC_50_ mortality at 0.451 µM OH•. Including DOC results in attenuation of toxicity to 6.5 µM OH•. At LC^50,^ the rate of hydroxyl radical generation in the 4.28 mg/L DOC exposures is ∼ 0.203 µM/h, while in the 0 mg/L DOC exposures, rate is ∼ 0.0141 µM/h. The rate difference is a result of much more TiO_2_ need to produce a toxic effect in suspensions with DOC. The toxic effect therefore appears to be a result of the rate of radical generation overwhelming the organism. In exposures with DOC, more radicals are required to cause induce equivalent toxic effects, a result that must be attributed to radical quenching by NOM.

Describing toxicity in terms of hydroxyl radicals may provide a more accurate description of toxicity than simply using TiO_2_ nanoparticle concentration. TiO_2_ alone is not toxic in the absence of UV-A light, exhibited by low mortality under no-UV or low UV conditions[14,15,24]. Rather, the hydroxyl radical is the actual toxicant in this relationship, and irradiated TiO_2_-mediated hydroxyl radical LC_50_ would be better described as such. Significantly more radicals are required to induce a similar mortality in suspensions with DOC, as compared to suspensions without DOC. Thus, longer exposure periods, and/or higher TiO_2_ concentrations in the system would be required for comparable toxicity. Researchers have previously shown that NOM provides protection from ROS from irradiated TiO_2_, but had not directly correlated changes in radical rate to toxicity[24]. These results demonstrate the important of including both irradiant intensity, and DOC in toxicity measurements, as these factors are integral in assessing toxicity.

## Acknowledgement

This work would not have been possible without, and is dedicated in memory of, Dr. Stephen J. Klaine

## Disclaimer

The authors have no conflicts of interests to disclose, financial or otherwise

