## Supplemental Information for "Quantitative measurements of hydroxyl radicals generated by irradiated titanium dioxide nanoparticle suspensions"

### NOM Characterization

*Table S1: Elemental Analysis of Suwanee River Water. Elemental analysis of filtered Suwannee River Water by ICP/MS. Content was consistent with elemental analysis of Suwannee River NOM certified by the Humic Substances Society.*

| Sample | P | K | Ca | Mg | Zn | Cu |
|---|---|---|---|---|---|---|
| | ppm | ppm | ppm | ppm | ppm | ppm |
| 1 | 0.071 | 1.735 | 2.698 | 1.186 | 0.012 | 0.008 |
| 2 | 0.066 | 1.699 | 2.669 | 1.184 | 0.012 | 0.009 |
| 3 | 0.074 | 1.683 | 2.673 | 1.191 | 0.012 | 0.007 |
| Average | 0.07 | 1.706 | 2.68 | 1.187 | 0.012 | 0.008 |
| Std Dev | 0.003 | 0.022 | 0.013 | 0.003 | 0 | 0.001 |
| Sample | Mn | Fe | S | Na | B | Al |
| | ppm | ppm | ppm | ppm | ppm | ppm |
| 1 | 0.03 | 1.15 | 0.923 | 5.647 | 0.025 | 0.83 |
| 2 | 0.029 | 1.143 | 0.937 | 5.536 | 0.024 | 0.804 |
| 3 | 0.028 | 1.168 | 0.916 | 5.505 | 0.024 | 0.834 |
| Average | 0.029 | 1.154 | 0.925 | 5.563 | 0.024 | 0.823 |
| Std Dev | 0.001 | 0.011 | 0.009 | 0.061 | 0.001 | 0.013 |

### Calibration curves for Hydroxyl Radical Quantitation

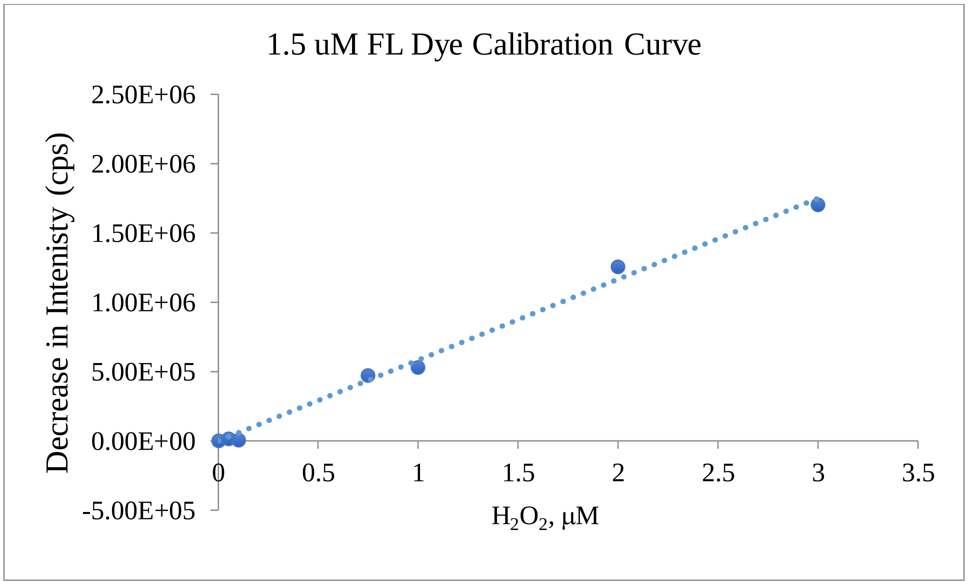


Figure S1: 1.5 µM fluorescein dye Hydroxyl Radical Concentration Calibration Curves produced by the catalytic cleavage of $H_2O_2$ by horseradish peroxidase. Measured by fluorescence Spectroscopy. ($R^2$=0.9963)

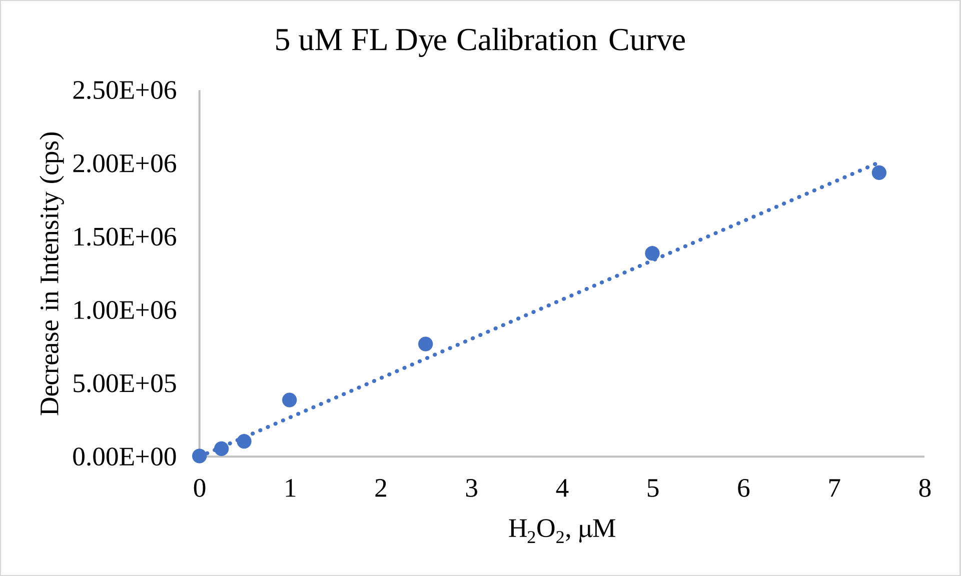


Figure S2: 5.0 µM fluorescein dye Hydroxyl Radical Concentration Calibration Curves produced by the catalytic cleavage of $H_2O_2$ by horseradish peroxidase. Measured by fluorescence Spectroscopy. ($R^2$=0.9905)

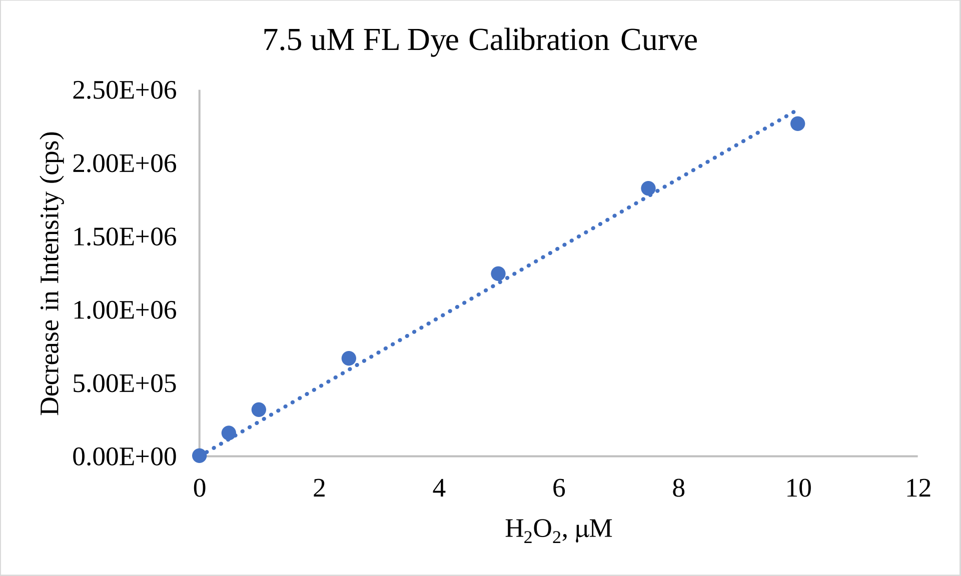


Figure S3: 7.5 µM fluorescein dye Hydroxyl Radical Concentration Calibration Curves produced by the catalytic cleavage of $H_2O_2$ by horseradish peroxidase. Measured by fluorescence Spectroscopy. ($R^2$=0.9935)

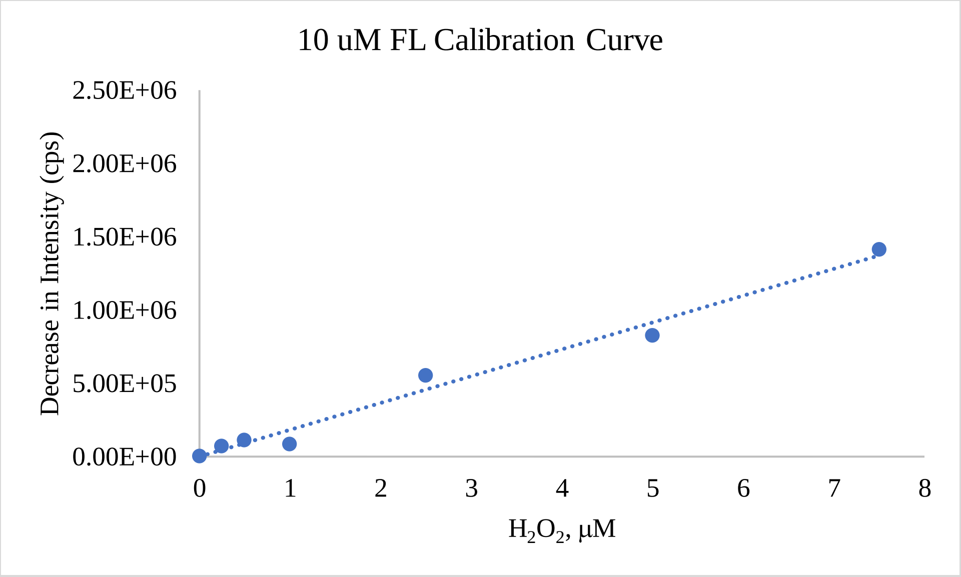


Figure S4: 10.0 µM fluorescein dye Hydroxyl Radical Concentration Calibration Curves produced by the catalytic cleavage of $H_2O_2$ by horseradish peroxidase. Measured by fluorescence Spectroscopy.($R_2$=0.9829)

### Lighting Characterization

The absolute irradiance of the light sources was measured periodically throughout testing to ensure no decrease in irradiant output occurred. Intensity was measured across the light spectrum**.** Dark conditions (no lights), exhibited no peaks within the UV spectra. UV irradiance was measured as average intensity across the UV-A spectrum (320 - 400 nm). The lowest UV intensity exposure measured an average irradiant intensity of 2.671 (±0.004) µW/$cm_2$/nm with maximum intensity at 365 nm measured 6.263 (±0.012) µW/$cm_2$/nm. Midrange intensity generated average irradiant intensity of 4.301 (±0.017) µW/$cm_2$/nm, with a maximum peak of 9.578 (±0.034) µW/$cm_2$/nm at 365 nm. The highest light intensity for exposures was measured at 5.188 (±0.032) µW/$cm_2$/nm with a maximum intensity peak of 10.833 (±0.054) µW/$cm_2$/nm at 365 nm. The light intensities were compared to peak intensity from the sun on a summer day in Pendleton, SC (34.6518 ° N, 82.7838°W). Average absolute irradiance was measured as 45.08

(±0.012) $\mu W/cm^2/nm$ and a maximum irradiance at 365 nm of 50.86 (±0.023) $\mu W/cm^2$. Output spectra shown in **Figure and Table.**

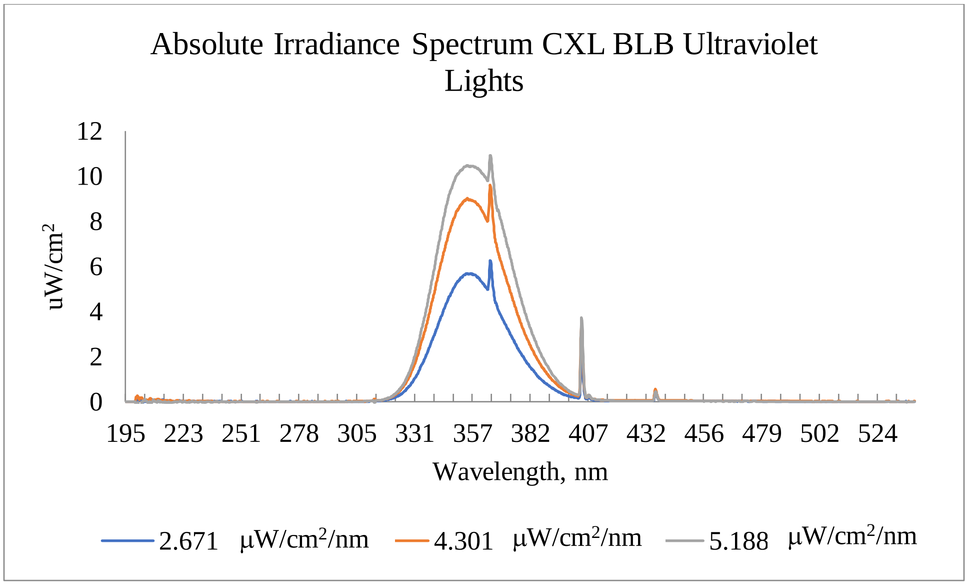


Figure S5: Absolute Irradiance Spectra of CLX-Blacklight Blue Fluorescent Bulbs. Absolute Irradiance Spectra from UV treatments. Peak wavelength is measured at 365 nm.

*Table S2. Absolute Irradiance from CLX-Blacklight Blue Fluorescent Bulb Ultraviolet-A (UV-A) Light Exposure System.*[a]

| Light | Maximum Absolute Irradiance (365 nm) | Average Absolute Irradiance (320 nm to 400 nm) |
|---|---|---|
| Dual Light | 10.833 ± 0.054 | 5.188 ± 0.032 |
| Single Light | 9.578 ± 0.034 | 4.301 ± 0.017 |
| Screened Light | 6.263 ± 0.012 | 2.671 ± 0.004 |
| Sunlight | 50.86 ± 0.023 | 45.08 ± 0.012 |

[a]Values are means ± standard deviation.

*Table S3. Decrease of UV-A Transmission through Moderately Hard Water with Increasing NOM Concentrations. Decrease in incident light intensity as DOC concentration is increased in MHW.*

| **% Transmission at 365 nm** | | | | | | | | | |
|---|---|---|---|---|---|---|---|---|---|
| 0 mg/L DOC | 0.5 mg/L DOC | 1.0 mg/L DOC | 2.5 mg/L DOC | 5.0 mg/L DOC | 10 mg/L DOC | 20 mg/L DOC | 40 mg/L DOC | 50 mg/L DOC | 60 mg/L DOC |
| **100.05** | **99.72** | **98.53** | **95.97** | **90.85** | **84.40** | **68.09** | **54.08** | **46.01** | **40.85** |

DOC=Dissolved organic carbon

Characterization of $TiO_2$ Nanoparticles

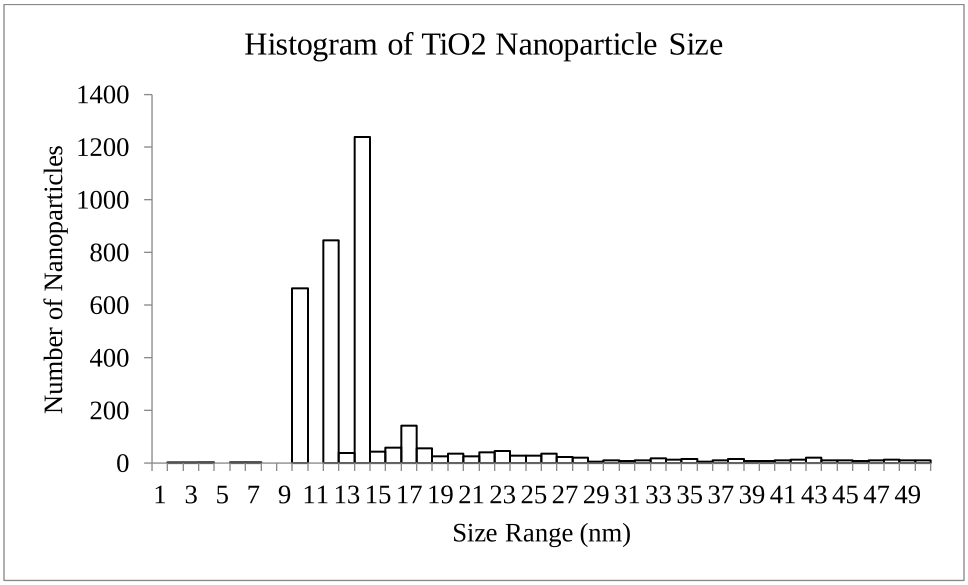


Figure S6. Size of $TiO_2$ Nanoparticle as Measured by Transmission Electron Microscope. Size of $TiO_2$ nanoparticles determined by TEM. Nanoparticle mean size: 21 ± 19 nm, Nanoparticle mode: 14 nm, N=4079.

Zeta Potential Analysis of Surface Charge

*Table S4: Zeta Potential of $TiO_2$/Natural Organic Matter (NOM) Suspension. 0 hour zeta potentials of suspensions of increasing $TiO_2$ concentrations and increasing DOC concentrations*

| | 0 mg/L DOC | 1.57 mg/L DOC | 2.95 mg/L DOC | 4.28 mg/L DOC | 5.71 mg/L DOC |
|---|---|---|---|---|---|
| $TiO_2$ mg/L | Zeta potential (mV) | Zeta potential (mV) | Zeta potential (mV) | Zeta potential (mV) | Zeta potential (mV) |
| 0 | n.a. | -11.43 ± 0.21 | -11.54 ± 2.66 | -14.07 ± 2.12 | -15.07 ± 0.94 |
| 0.5 | -13.97 ± 0.58 | -17.67 ± 0.9 | -15.63 ± 0.35 | -14.37 ± 0.51 | -16.93 ± 1.58 |
| 1 | -14.83 ± 0.59 | -16.03 ± 0.32 | -15.8 ± 0.69 | -12.83 ± 0.65 | -14.87 ± 0.1.57 |
| 3.5 | -14.03 ± 0.12 | -13.1 ± 0.27 | -15.67 ± 1.05 | -15.9 ± 0.36 | -16.27 ± 1.01 |
| 5 | -13.6 ± 0.56 | -17.2 ± 0.58 | -16.9 ± 0.4 | -20.37 ± 0.38 | -15.33 ± 1.29 |
| 7 | -14.33 ± 0.55 | -15.77 ± 0.58 | -16.9 ± 0.61 | -16.57 ± 0.55 | -16.5 ± 0.53 |
| 10.5 | -14.27 ± 0.23 | -15.57 ± 0.51 | -15.13 ± 0.12 | -14.93 ± 0.31 | -20.03 ± 0.38 |
| 14 | -14.37 ± 0.0.28 | -16.3 ± 0.82 | -12.1 ± 0.36 | -15.57 ± 0.21 | -15.97 ± 0.25 |

[a]Values are means ± standard deviation
$TiO_2$=titanium dioxide, DOC=dissolved organic carbon

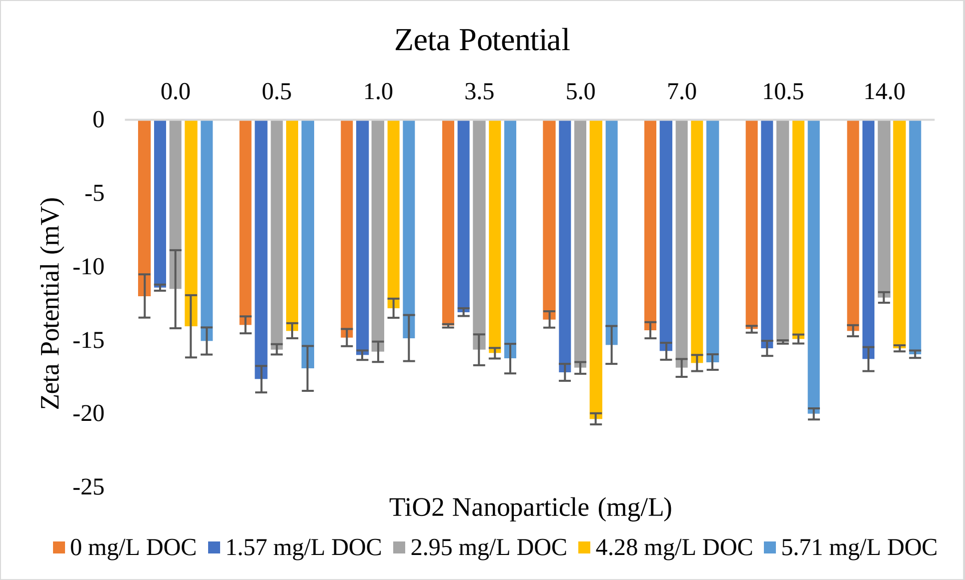


Figure S7: Change in Zeta Potential with Increasing $TiO_2$ and DOC Concentrations. Zeta potential of suspensions on increasing DOC and $TiO_2$ concentrations

*DLS Measurement of Z-average $TiO_2$ aggregate sizes*

Categorically organized by DOC across 48 hours, there was no statistically significant differences between the group means as determined by a one-way ANOVA: $DOC_0$:($F$(2,81) = 0.8252, $p$= 0.4418), $DOC_{[1.57]}$ :($F$(2,81) = 1.17958, $p$= 0.1725), $DOC_{[2.95]}$:($F$(2,81)= 1.4391, $p$= 0.2431), $DOC_{[4.28]}$ : ($F$(2,80) = 0.5983, $p$= 0.5522), $DOC_{[5.71]}$:($F$(2,81) = 0.0866, $p$= 0.9171). These results are shown in **TABLE S5 to S9.**

*Table S5: ANOVA analysis of z-average $TiO_2$ aggregate size with 0.5 mg/L DOC across 48 hours, as measured by Dynamic Light Scattering.*

| Analysis of Variance: 0.5 mg/L DOC | | | | | |
|---|---|---|---|---|---|
| **Source** | **DF** | **Sum of Squares** | **Mean Square** | **F Ratio** | **Prob > F** |
| Time | 2 | 16705.45 | 8352.7 | 0.8252 | 0.4418 |
| Error | 81 | 819919.25 | 10122.5 | | |
| C. Total | 83 | 836624.70 | | | |
| **Means for One-way ANOVA** | | | | | |
| **Hour** | **Number** | **Mean** | **Std Error** | **Lower 95%** | **Upper 95%** |
| 0 | 28 | 196.071 | 19.014 | 158.24 | 233.9 |
| 24 | 28 | 189.643 | 19.014 | 151.81 | 227.47 |
| 48 | 28 | 163.464 | 19.014 | 125.63 | 201.3 |

*Std Error used a pooled estimate for error variance

*Table S6: ANOVA analysis of z-average $TiO_2$ aggregate size with 1.57 mg/L DOC across 48 hours, as measured by Dynamic Light Scattering.*

| Analysis of Variance: 1.57 mg/L DOC | | | | | |
|---|---|---|---|---|---|
| **Source** | **DF** | **Sum of Squares** | **Mean Square** | **F Ratio** | **Prob > F** |
| Time | 2 | 26402.00 | 13201.00 | 1.7958 | 0.1725 |
| Error | 81 | 595436.32 | 7351.10 | | |
| C. Total | 83 | 621838.32 | | | |
| **Means for One-way ANOVA** | | | | | |
| **Hour** | **Number** | **Mean** | **Std Error** | **Lower 95%** | **Upper 95%** |
| 0 | 28 | 190.54 | 16.203 | 158.30 | 222.77 |
| 24 | 28 | 190.18 | 16.203 | 157.94 | 222.42 |
| 48 | 28 | 152.75 | 16.203 | 120.51 | 184.99 |

*Std Error used a pooled estimate for error variance

*Table S7: ANOVA analysis of z-average $TiO_2$ aggregate size with 2.95 mg/L DOC across 48 hours, as measured by Dynamic Light Scattering.*

| Analysis of Variance: 2.95 mg/L DOC | | | | | |
|---|---|---|---|---|---|
| **Source** | **DF** | **Sum of Squares** | **Mean Square** | **F Ratio** | **Prob > F** |
| Time | 2 | 33047.21 | 16523.6 | 1.4391 | 0.2431 |
| Error | 81 | 930010.93 | 11481.6 | | |
| C. Total | 83 | 963058.14 | | | |
| **Means for One-way ANOVA** | | | | | |
| **Hour** | **Number** | **Mean** | **Std Error** | **Lower 95%** | **Upper 95%** |
| 0 | 28 | 184.036 | 20.25 | 143.74 | 224.33 |
| 24 | 28 | 215.679 | 20.25 | 175.39 | 255.97 |
| 48 | 28 | 167.929 | 20.25 | 127.64 | 208.22 |

*Std Error used a pooled estimate for error variance

*Table S8: ANOVA analysis of z-average $TiO_2$ aggregate size with 4.28 mg/L DOC across 48 hours, as measured by Dynamic Light Scattering*.

| Analysis of Variance: 4.28 mg/L DOC | | | | | |
|---|---|---|---|---|---|
| **Source** | **DF** | **Sum of Squares** | **Mean Square** | **F Ratio** | **Prob > F** |
| Time | 2 | 5165.56 | 2582.78 | 0.5983 | 0.5522 |
| Error | 80 | 345375.31 | 4317.19 | | |
| C. Total | 82 | 350540.87 | | | |
| **Means for One-way ANOVA** | | | | | |
| **Hour** | **Number** | **Mean** | **Std Error** | **Lower 95%** | **Upper 95%** |
| 0 | 27 | 174.704 | 12.645 | 149.54 | 199.87 |
| 24 | 28 | 157.536 | 12.417 | 132.82 | 182.25 |
| 48 | 28 | 158.214 | 12.417 | 133.5 | 182.93 |

*Std Error used a pooled estimate for error variance

*Table S9: ANOVA analysis of z-average $TiO_2$ aggregate size with 5.71 mg/L DOC across 48 hours, as measured by Dynamic Light Scattering*.

| Analysis of Variance: 5.71 mg/L DOC | | | | | |
|---|---|---|---|---|---|
| **Source** | **DF** | **Sum of Squares** | **Mean Square** | **F Ratio** | **Prob > F** |
| Time | 2 | 1175.04 | 587.52 | 0.0866 | 0.9171 |
| Error | 80 | 542452.63 | 6780.66 | | |
| C. Total | 82 | 543627.66 | | | |
| **Means for One-way ANOVA** | | | | | |
| **Hour** | **Number** | **Mean** | **Std Error** | **Lower 95%** | **Upper 95%** |
| 0 | 28 | 175.143 | 15.562 | 144.17 | 206.11 |
| 24 | 27 | 181.593 | 15.847 | 150.06 | 213.13 |
| 48 | 28 | 172.607 | 15.562 | 141.64 | 203.58 |

*Std Error used a pooled estimate for error variance

Considering the effect that $TiO_2$ had on the group mean of z-average aggregate size at each timepoint, there were no statistically significant differences at the 0 hour timepoint: ($F(4,136) = 0.4989$, $p = 0.7396$), the 24 hour time point: ($F(4,136) = 1.0699$, $p = 0.3740$), the 48 hour time point: ($F(6,136) = 2208$, $p = 0.9265$). These results are demonstrated in **TABLE S10 to S12.**

*Table S10: ANOVA analysis of z-average $TiO_2$ aggregate size at 0-hour as DOC (mg/L) is increased, as measured by Dynamic Light Scattering.*

| Analysis of Variance: DOC at 0-hr | | | | | |
|---|---|---|---|---|---|
| **Source** | **DF** | **Sum of Squares** | **Mean Square** | **F Ratio** | **Prob > F** |
| DOC | 4 | 9802.35 | 2450.59 | 0.4948 | 0.7396 |
| Error | 134 | 663636.84 | 4952.51 | | |
| C. Total | 138 | 673439.19 | | | |
| **Means for One-way ANOVA: DOC at 0-hr** | | | | | |
| **DOC Conc** | **Number** | **Mean** | **Std Error** | **Lower 95%** | **Upper 95%** |
| 0 | 28 | 196.071 | 13.299 | 169.77 | 222.38 |
| 1.57 | 28 | 190.536 | 13.299 | 164.23 | 216.84 |
| 2.95 | 28 | 184.036 | 13.299 | 157.73 | 210.34 |
| 4.28 | 27 | 174.704 | 13.544 | 147.92 | 201.49 |
| 5.71 | 28 | 175.143 | 13.299 | 148.84 | 201.45 |

*Std Error used a pooled estimate for error variance

*Table S10: ANOVA analysis of z-average $TiO_2$ aggregate size at 24-hour as DOC (mg/L) is increased, as measured by Dynamic Light Scattering.*

| Analysis of Variance: DOC at 24-hr | | | | | |
|---|---|---|---|---|---|
| **Source** | **DF** | **Sum of Squares** | **Mean Square** | **F Ratio** | **Prob > F** |
| DOC | 4 | 48604.7 | 12151.2 | 1.0699 | 0.374 |
| Error | 134 | 1521912.1 | 11357.6 | | |
| C. Total | 138 | 1570516.8 | | | |
| **Means for One-way ANOVA: DOC at 24-hr** | | | | | |
| **DOC Conc** | **Number** | **Mean** | **Std Error** | **Lower 95%** | **Upper 95%** |
| 0 | 28 | 189.643 | 20.14 | 149.81 | 229.48 |
| 1.57 | 28 | 190.179 | 20.14 | 150.34 | 230.01 |
| 2.95 | 28 | 215.679 | 20.14 | 175.84 | 255.51 |
| 4.28 | 28 | 157.536 | 20.14 | 117.7 | 197.37 |
| 5.71 | 27 | 181.593 | 20.51 | 141.03 | 222.16 |

*Std Error used a pooled estimate for error variance

*Table S10: ANOVA analysis of z-average $TiO_2$ aggregate size at 48-hour as DOC (mg/L) is increased, as measured by Dynamic Light Scattering.*

| **Analysis of Variance: DOC at 48-hr** | | | | | |
|---|---|---|---|---|---|
| **Source** | **DF** | **Sum of Squares** | **Mean Square** | **F Ratio** | **Prob > F** |
| DOC | 4 | 6853.5 | 1713.38 | 0.2208 | 0.9265 |
| Error | 135 | 1047645.5 | 7760.34 | | |
| C. Total | 139 | 1054499 | | | |
| **Means for One-way ANOVA: DOC at 48-hr** | | | | | |
| **DOC Conc** | **Number** | **Mean** | **Std Error** | **Lower 95%** | **Upper 95%** |
| 0 | 28 | 163.464 | 16.648 | 130.54 | 196.39 |
| 1.57 | 28 | 152.75 | 16.648 | 119.83 | 185.67 |
| 2.95 | 28 | 167.929 | 16.648 | 135 | 200.85 |
| 4.28 | 28 | 158.214 | 16.648 | 125.29 | 191.14 |
| 5.71 | 28 | 172.607 | 16.648 | 139.68 | 205.53 |

*Std Error used a pooled estimate for error variance

Categorically organized by Light Intensity ($UV_i$) across 48 hours, there was no statistically significant differences between the group means as determined by a one-way ANOVA for $UV_{i[2.671]}$: ($F(2,102) = 1.290$, $p= 0.2797$)), $UV_{i[4.301]}$: ($F(2,102) = 1.315$, $p= 0.2729$)), and $UV_{i[5.177]}$: ($F(2,102) = 0.7053$, $p= 0.4964$)). There was a statistically significant difference between the group means as determined by one-way ANOVA $UV_{i\,0}$: ($F(2,102) = 5.8774$, $p= 0.0038$). A *post hoc* Tukey-Kramer HSD was run to determine levels of significance across 48 hours. These results are shown in **Table S11 to S14.**

*Table S11: ANOVA and Tukey-Kramer HSD* post hoc *analysis of z-average $TiO_2$ aggregate size under 0 μW/cm2/nm UV-A intensity across 48 hours, as measured by Dynamic Light Scattering.*

| Analysis of Variance: 0 μW/cm2/nm Light Intensity | | | | | |
|---|---|---|---|---|---|
| **Source** | **DF** | **Sum of Squares** | **Mean Square** | **F Ratio** | **Prob > F** |
| Time | 2 | 42707.37 | 21353.7 | 5.8774 | 0.0038* |
| Error | 102 | 370585.26 | 3633.2 | | |
| C. Total | 104 | 413292.63 | | | |
| **Means for One-way ANOVA** | | | | | |
| **Level** | **Number** | **Mean** | **Std Error** | **Lower 95%** | **Upper 95%** |
| 0 | 35 | 207.543 | 10.188 | 187.33 | 227.75 |
| 24 | 35 | 171.057 | 10.188 | 150.85 | 191.27 |
| 48 | 35 | 160.457 | 10.188 | 140.25 | 180.67 |
| **Tukey-Kramer HSD: 0 μW/cm2/nm Light Intensity** | | | | | |
| **q*** | **Alpha** | **Level** | | **Mean** | **Std Error** |
| 2.37843 | 0.05 | 0 | A | 207.54286 | 10.188 |
| | | 24 | B | 171.05714 | 10.188 |
| | | 48 | B | 160.45714 | 10.188 |

*Levels not connected by the same letter are significantly different
*Std Error used a pooled estimate for error variance

*Table S12: ANVOA analysis of z-average $TiO_2$ aggregate size under 2.671 μW/cm2/nm UV-A intensity across 48 hours, as measured by Dynamic Light Scattering.*

| Analysis of Variance: 2.671 μW/cm2/nm Light Intensity | | | | | |
|---|---|---|---|---|---|
| **Source** | **DF** | **Sum of Squares** | **Mean Square** | **F Ratio** | **Prob > F** |
| Time | 2 | 10569.77 | 5284.89 | 1.2901 | 0.2797 |
| Error | 102 | 417851.14 | 4096.58 | | |
| C. Total | 104 | 428420.91 | | | |
| **Means for One-way ANOVA** | | | | | |
| **Level** | **Number** | **Mean** | **Std Error** | **Lower 95%** | **Upper 95%** |
| 0 | 35 | 173 | 10.819 | 151.54 | 194.46 |
| 24 | 35 | 176.686 | 10.819 | 155.23 | 198.14 |
| 48 | 35 | 153.8 | 10.819 | 132.34 | 175.26 |

*Std Error used a pooled estimate for error variance

*Table S13: ANOVA analysis of z-average $TiO_2$ aggregate size under 4.301 μW/cm2/nm UV-A intensity across 48 hours, as measured by Dynamic Light Scattering.*

| **Analysis of Variance: 4.301 μW/cm2/nm Light Intensity** | | | | | |
|---|---|---|---|---|---|
| **Source** | **DF** | **Sum of Squares** | **Mean Square** | **F Ratio** | **Prob > F** |
| Time | 2 | 27467.2 | 13733.6 | 1.3153 | 0.2729 |
| Error | 102 | 1065063.7 | 10441.8 | | |
| C. Total | 104 | 1092530.9 | | | |
| **Means for One-way ANOVA** | | | | | |
| **Level** | **Number** | **Mean** | **Std Error** | **Lower 95%** | **Upper 95%** |
| 0 | 35 | 163.743 | 17.272 | 129.48 | 198 |
| 24 | 35 | 202.714 | 17.272 | 168.45 | 236.97 |
| 48 | 35 | 177.057 | 17.272 | 142.8 | 211.32 |

*Std Error used a pooled estimate for error variance

*Table S14: ANOVA analysis of z-average $TiO_2$ aggregate size under 5.177 μW/cm2/nm UV-A intensity across 48 hours, as measured by Dynamic Light Scattering.*

| **Analysis of Variance: 5.177 μW/cm2/nm Light Intensity** | | | | | |
|---|---|---|---|---|---|
| **Source** | **DF** | **Sum of Squares** | **Mean Square** | **F Ratio** | **Prob > F** |
| Time | 2 | 19954.3 | 9977.2 | 0.7053 | 0.4964 |
| Error | 102 | 1442993.3 | 14147 | | |
| C. Total | 104 | 1462947.7 | | | |
| **Means for One-way ANOVA** | | | | | |
| **Level** | **Number** | **Mean** | **Std Error** | **Lower 95%** | **Upper 95%** |
| 0 | 35 | 187.114 | 20.105 | 147.24 | 226.99 |
| 24 | 35 | 192.057 | 20.105 | 152.18 | 231.93 |
| 48 | 35 | 160.657 | 20.105 | 120.78 | 200.53 |

*Std Error used a pooled estimate for error variance

Considering the effect that $TiO_2$ had on the group mean of z-average aggregate size at each timepoint, there were no statistically significant differences at the 0 hour timepoint: ($F(3,135) = 2.2506$, $p = 0.0853$), the 24 hour time point: ($F(3,135) = 0.5606$, $p = 0.6419$), the 48 hour time point: ($F(3,136) = 0.4470$, $p = 0.7198$). These results are demonstrated in **TABLE S15 to S17.**

*Table S15: ANOVA analysis of z-average $TiO_2$ aggregate size comparing differences based on light intensities at 0-hour, as measured by Dynamic Light Scattering.*

| Analysis of Variance: Light Intensity at 0-hour | | | | | |
|---|---|---|---|---|---|
| **Source** | **DF** | **Sum of Squares** | **Mean Square** | **F Ratio** | **Prob > F** |
| Light | 3 | 32076.58 | 10692.2 | 2.2506 | 0.0853 |
| Error | 135 | 641362.61 | 4750.8 | | |
| C. Total | 138 | 673439.19 | | | |
| **Means for One-way ANOVA** | | | | | |
| **Light Intensity** | **Number** | **Mean** | **Std Error** | **Lower 95%** | **Upper 95%** |
| 0 | 35 | 207.54 | 11.651 | 184.50 | 230.58 |
| 2.671 | 35 | 173.00 | 11.651 | 149.96 | 196.04 |
| 4.301 | 34 | 168.56 | 11.821 | 145.18 | 191.94 |
| 5.177 | 35 | 187.11 | 11.651 | 164.07 | 210.16 |

*Std Error used a pooled estimate for error variance

*Table S16: ANOVA analysis of z-average $TiO_2$ aggregate size comparing differences based on light intensities at 24-hour, as measured by Dynamic Light Scattering.*

| Analysis of Variance: Light Intensity at 24-hour | | | | | |
|---|---|---|---|---|---|
| **Source** | **DF** | **Sum of Squares** | **Mean Square** | **F Ratio** | **Prob > F** |
| Light | 3 | 19324.4 | 6441.5 | 0.5606 | 0.6419 |
| Error | 135 | 1551192.4 | 11490.3 | | |
| C. Total | 138 | 1570516.8 | | | |
| **Means for One-way ANOVA** | | | | | |
| **Light Intensity** | **Number** | **Mean** | **Std Error** | **Lower 95%** | **Upper 95%** |
| 0 | 35 | 171.057 | 18.119 | 135.22 | 206.89 |
| 2.671 | 34 | 181.882 | 18.383 | 145.53 | 218.24 |
| 4.301 | 35 | 202.714 | 18.119 | 166.88 | 238.55 |
| 5.177 | 35 | 192.057 | 18.119 | 156.22 | 227.89 |

*Std Error used a pooled estimate for error variance

*Table S17: ANOVA analysis of z-average $TiO_2$ aggregate size comparing differences based on light intensities at 48-hour, as measured by Dynamic Light Scattering.*

| Analysis of Variance: Light Intensity at 48-hour | | | | | |
|---|---|---|---|---|---|
| **Source** | **DF** | **Sum of Squares** | **Mean Square** | **F Ratio** | **Prob > F** |
| Light | 3 | 10296.9 | 3432.31 | 0.447 | 0.7198 |
| Error | 136 | 1044202.1 | 7677.96 | | |
| C. Total | 139 | 1054499 | | | |
| **Means for One-way ANOVA** | | | | | |
| **Light Intensity** | **Number** | **Mean** | **Std Error** | **Lower 95%** | **Upper 95%** |
| 0 | 35 | 160.457 | 14.811 | 131.17 | 189.75 |
| 2.671 | 35 | 153.8 | 14.811 | 124.51 | 183.09 |
| 4.301 | 35 | 177.057 | 14.811 | 147.77 | 206.35 |
| 5.177 | 35 | 160.657 | 14.811 | 131.37 | 189.95 |

*Std Error used a pooled estimate for error variance

Categorically organized by $TiO_2$ concentration across 48 hours, there were no statistically significant differences between group means as determined by one-way ANOVA for $TiO_{2\,[0.5]}$: ($F$(2,57) = 0.1988, $p$= 0.8203)), $TiO_{2\,[1.0]}$: ($F$(2,57) = 0.8540, $p$= 0.4311)), $TiO_{2\,[5.0]}$: ($F$(2,57) = 0.1988, $p$= 0.8203)),$TiO_{2\,[7.0]}$ ($F$(2,57) = 2.0982, $p$= 0.1320)), $TiO_{2\,[10.5]}$ ($F$(2,57) = 0.4926, $p$= 0.6136)). There were statistically significant differences between group means as determined by one-way ANOVA for $TiO_{2\,[3.5]}$: ($F$(2,57) = 6.1514, $p$= 0.0038)),and $TiO_{2\,[14.0]}$: ($F$(2,57) = 3.9067, $p$= 0.0257)). A *post hoc* Tukey-Kramer HSD was run to determine levels of significance across 48 hours. These results are shown in **Table S18 to S24.**

*Table S18: ANOVA analysis of z-average $TiO_2$ aggregate size with 0.5 mg/L $TiO_2$ across 48 hours, as measured by Dynamic Light Scattering.*

| Analysis of Variance: 0.5 mg/L TiO2 | | | | | |
|---|---|---|---|---|---|
| **Source** | **DF** | **Sum of Squares** | **Mean Square** | **F Ratio** | **Prob > F** |
| Time | 2 | 955.73 | 477.87 | 0.1988 | 0.8203 |
| Error | 57 | 137036.45 | 2404.15 | | |
| C. Total | 59 | 137992.18 | | | |
| **Means for One-way ANOVA** | | | | | |
| **Hour** | **Number** | **Mean** | **Std Error** | **Lower 95%** | **Upper 95%** |
| 0 | 20 | 115.45 | 10.964 | 93.495 | 137.4 |
| 24 | 20 | 118.65 | 10.964 | 96.695 | 140.6 |
| 48 | 20 | 109.05 | 10.964 | 87.095 | 131.0 |

*Std Error used a pooled estimate for error variance

*Table S19: ANOVA analysis of z-average $TiO_2$ aggregate size with 1.0 mg/L $TiO_2$ across 48 hours, as measured by Dynamic Light Scattering.*

| Analysis of Variance: 1.0 mg/L TiO2 | | | | | |
|---|---|---|---|---|---|
| **Source** | **DF** | **Sum of Squares** | **Mean Square** | **F Ratio** | **Prob > F** |
| Time | 2 | 5031.3 | 2515.65 | 0.854 | 0.4311 |
| Error | 57 | 167901.7 | 2945.64 | | |
| C. Total | 59 | 172933 | | | |
| **Means for One-way ANOVA** | | | | | |
| **Hour** | **Number** | **Mean** | **Std Error** | **Lower 95%** | **Upper 95%** |
| 0 | 20 | 107.95 | 12.136 | 83.65 | 132.25 |
| 24 | 20 | 127.45 | 12.136 | 103.15 | 151.75 |
| 48 | 20 | 108.1 | 12.136 | 83.8 | 132.4 |

*Std Error used a pooled estimate for error variance

*Table S20: ANOVA and associated Tukey-Kramer HSD* post hoc *analysis of z-average $TiO_2$ aggregate size with 3.5 mg/L $TiO_2$ across 48 hours, as measured by Dynamic Light Scattering.*

| Analysis of Variance: 3.5 mg/L TiO2 | | | | | |
|---|---|---|---|---|---|
| **Source** | **DF** | **Sum of Squares** | **Mean Square** | **F Ratio** | **Prob > F** |
| Time | 2 | 39265.56 | 19632.8 | 6.1514 | 0.0038* |
| Error | 56 | 178730.17 | 3191.6 | | |
| C. Total | 58 | 217995.73 | | | |
| **Means for One-way ANOVA** | | | | | |
| **Hour** | **Number** | **Mean** | **Std Error** | **Lower 95%** | **Upper 95%** |
| 0 | 20 | 167.45 | 12.633 | 142.14 | 192.76 |
| 24 | 19 | 149.368 | 12.961 | 123.41 | 175.33 |
| 48 | 20 | 106.4 | 12.633 | 81.09 | 131.71 |
| **Tukey-Kramer HSD for 3.5 mg/L TiO2** | | | | | |
| **q*** | **Alpha** | **Level** | | **Mean** | **Std Error** |
| 2.40757 | 0.05 | 0 | A | 167.45 | 12.633 |
| | | 24 | AB | 149.367 | 12.961 |
| | | 48 | B | 106.4 | 12.633 |

*Levels not connected by the same letter are significantly different
*Std Error used a pooled estimate for error variance

*Table S21: ANOVA analysis of z-average $TiO_2$ aggregate size with 5.0 mg/L $TiO_2$ across 48 hours, as measured by Dynamic Light Scattering.*

| Analysis of Variance: 5.0 mg/L TiO2 | | | | | |
|---|---|---|---|---|---|
| **Source** | **DF** | **Sum of Squares** | **Mean Square** | **F Ratio** | **Prob > F** |
| Time | 2 | 18885.7 | 9442.85 | 1.4784 | 0.2368 |
| Error | 56 | 357687.82 | 6387.28 | | |
| C. Total | 58 | 376573.53 | | | |
| **Means for One-way ANOVA** | | | | | |
| **Hour** | **Number** | **Mean** | **Std Error** | **Lower 95%** | **Upper 95%** |
| 0 | 19 | 191.368 | 18.335 | 154.64 | 228.1 |
| 24 | 20 | 178.8 | 17.871 | 143.00 | 214.6 |
| 48 | 20 | 148.7 | 17.871 | 112.90 | 184.5 |

*Std Error used a pooled estimate for error variance

*Table S22: ANOVA analysis of z-average $TiO_2$ aggregate size with 7.0 mg/L $TiO_2$ across 48 hours, as measured by Dynamic Light Scattering.*

| Analysis of Variance: 7.0 mg/L TiO2 | | | | | |
|---|---|---|---|---|---|
| **Source** | **DF** | **Sum of Squares** | **Mean Square** | **F Ratio** | **Prob > F** |
| Time | 2 | 11688.13 | 5844.07 | 2.0982 | 0.132 |
| Error | 57 | 158758.85 | 2785.24 | | |
| C. Total | 59 | 170446.98 | | | |
| **Means for One-way ANOVA** | | | | | |
| **Hour** | **Number** | **Mean** | **Std Error** | **Lower 95%** | **Upper 95%** |
| 0 | 20 | 215.05 | 11.801 | 191.42 | 238.68 |
| 24 | 20 | 187.95 | 11.801 | 164.32 | 211.58 |
| 48 | 20 | 183.45 | 11.801 | 159.82 | 207.08 |

*Std Error used a pooled estimate for error variance

*Table S23: ANOVA analysis of z-average $TiO_2$ aggregate size with 10.5 mg/L $TiO_2$ across 48 hours, as measured by Dynamic Light Scattering*.

| Analysis of Variance: 10.5 mg/L TiO2 | | | | | |
|---|---|---|---|---|---|
| **Source** | **DF** | **Sum of Squares** | **Mean Square** | **F Ratio** | **Prob > F** |
| Time | 2 | 5433.1 | 2716.55 | 0.4926 | 0.6136 |
| Error | 57 | 314327.9 | 5514.52 | | |
| C. Total | 59 | 319761 | | | |
| **Means for One-way ANOVA** | | | | | |
| **Hour** | **Number** | **Mean** | **Std Error** | **Lower 95%** | **Upper 95%** |
| 0 | 20 | 235.8 | 16.605 | 202.55 | 269.05 |
| 24 | 20 | 222.25 | 16.605 | 189 | 255.5 |
| 48 | 20 | 245.45 | 16.605 | 212.2 | 278.7 |

*Std Error used a pooled estimate for error variance

*Table S24: ANOVA and Tukey-Kramer HSD* post *hoc analysis of z-average $TiO_2$ aggregate size with 14.0 mg/L $TiO_2$ across 48 hours, as measured by Dynamic Light Scattering.*

| Analysis of Variance: 14.0 mg/L TiO2 | | | | | |
|---|---|---|---|---|---|
| **Source** | **DF** | **Sum of Squares** | **Mean Square** | **F Ratio** | **Prob > F** |
| Time | 2 | 76329.23 | 38164.6 | 3.9067 | 0.0257* |
| Error | 57 | 556840.95 | 9769.1 | | |
| C. Total | 59 | 633170.18 | | | |
| **Means for One-way ANOVA** | | | | | |
| **Hour** | **Number** | **Mean** | **Std Error** | **Lower 95%** | **Upper 95%** |
| 0 | 20 | 256.45 | 22.101 | 212.19 | 300.71 |
| 24 | 20 | 322.4 | 22.101 | 278.14 | 366.66 |
| 48 | 20 | 239.8 | 22.101 | 195.54 | 284.06 |
| **Tukey-Kramer HSD for 3.5 mg/L TiO2** | | | | | |
| **q*** | **Alpha** | **Level** | | **Mean** | **Std Error** |
| 2.40642 | 0.05 | 24 | A | 322.4 | 22.101 |
| | | 0 | AB | 256.45 | 22.101 |
| | | 48 | B | 239.8 | 22.101 |

*Levels not connected by the same letter are significantly different
*Std Error used a pooled estimate for error variance

Considering the effect that $TiO_2$ had on the group mean of z-average aggregate size at each timepoint, there were statistically significant differences at the 0 hour timepoint: ($F$(6,132) = 31.0357, $p$ = <0.0001), the 24 hour time point: ($F$(6,132) = 13.0286, $p$ = <0.0001), the 48 hour time point: ($F$(6,132) = 16.4337, $p$ = <0.0001). A *post hoc* Tukey-Kramer HSD was run to determine levels of significance between the concentrations of $TiO_2$ at each of these timepoints. These results are shown in **Table S25 to S27.**

*Table S25: ANOVA and Tukey-Kramer HSD* post *hoc analysis of z-average $TiO_2$ aggregate size at 0 hour as $TiO_2$ concentration was increased, measured by Dynamic Light Scattering.*

| **Analysis of Variance: $TiO_2$ at 0-hour** | | | | | |
|---|---|---|---|---|---|
| **Source** | **DF** | **Sum of Squares** | **Mean Square** | **F Ratio** | **Prob > F** |
| $TiO_2$ | 6 | 394086.82 | 65681.1 | 31.0357 | <.0001* |
| Error | 132 | 279352.37 | 2116.3 | | |
| C. Total | 138 | 673439.19 | | | |

| **Means for One-way ANOVA** | | | | | |
|---|---|---|---|---|---|
| **Level** | **Number** | **Mean** | **Std Error** | **Lower 95%** | **Upper 95%** |
| 0.5 | 20 | 115.45 | 10.287 | 95.1 | 135.8 |
| 1.0 | 20 | 107.95 | 10.287 | 87.6 | 128.3 |
| 3.5 | 20 | 167.45 | 10.287 | 147.1 | 187.8 |
| 5.0 | 19 | 191.37 | 10.554 | 170.5 | 212.3 |
| 7.0 | 20 | 215.05 | 10.287 | 194.7 | 235.4 |
| 10.5 | 20 | 235.80 | 10.287 | 215.5 | 256.2 |
| 14.0 | 20 | 256.45 | 10.287 | 236.1 | 276.8 |

| **Tukey-Kramer HSD: $TiO_2$ at 0-hour** | |
|---|---|
| **q*** | **Alpha** |
| 2.99433 | 0.05 |

| **Connecting Letters Report** | | | |
|---|---|---|---|
| **Level** | | **Mean** | **Std Error** |
| 14.0 | A | 256.45 | 10.287 |
| 10.5 | A | 235.80 | 10.287 |
| 7.0 | AB | 215.05 | 10.287 |
| 5.0 | BC | 191.37 | 10.554 |
| 3.5 | C | 167.45 | 10.287 |
| 1.0 | D | 107.95 | 10.287 |
| 0.5 | D | 115.45 | 10.287 |

*Levels not connected by the same letter are significantly different
*Std Error used a pooled estimate for error variance

*Table S26: ANOVA and Tukey-Kramer HSD* post *hoc analysis of z-average $TiO_2$ aggregate size at 24-hour as $TiO_2$ concentration was increased, measured by Dynamic Light Scattering.*

| Analysis of Variance: $TiO_2$ at 24-hour | | | | | |
|---|---|---|---|---|---|
| **Source** | **DF** | **Sum of Squares** | **Mean Square** | **F Ratio** | **Prob > F** |
| TiO2 | 6 | 584142.2 | 97357 | 13.0286 | <.0001* |
| Error | 132 | 986374.6 | 7472.5 | | |
| C. Total | 138 | 1570516.8 | | | |

| Means for One-way ANOVA | | | | | |
|---|---|---|---|---|---|
| **Level** | **Number** | **Mean** | **Std Error** | **Lower 95%** | **Upper 95%** |
| 0.5 | 20 | 118.65 | 19.329 | 80.41 | 156.89 |
| 1 | 20 | 127.45 | 19.329 | 89.21 | 165.69 |
| 3.5 | 19 | 149.37 | 19.832 | 110.14 | 188.60 |
| 5 | 20 | 178.80 | 19.329 | 140.56 | 217.04 |
| 7 | 20 | 187.95 | 19.329 | 149.71 | 226.19 |
| 10.5 | 20 | 222.25 | 19.329 | 184.01 | 260.49 |
| 14 | 20 | 322.40 | 19.329 | 284.16 | 360.64 |

| Tukey Kramer HSD: $TiO_2$ at 24-hour | |
|---|---|
| **q*** | **Alpha** |
| 2.99433 | 0.05 |

| Connecting Letters Report | | | |
|---|---|---|---|
| **Level** | | **Mean** | **Std Error** |
| 14 | A | 322.4 | 19.329 |
| 10.5 | B | 222.25 | 19.329 |
| 7 | BC | 187.95 | 19.329 |
| 5 | BC | 178.8 | 19.329 |
| 3.5 | BC | 149.37 | 19.832 |
| 1 | C | 127.45 | 19.329 |
| 0.5 | C | 118.65 | 19.329 |

*Levels not connected by the same letter are significantly different
*Std Error used a pooled estimate for error variance

*Table S27: ANOVA and Tukey-Kramer HSD* post *hoc analysis of z-average $TiO_2$ aggregate size at 48-hour as $TiO_2$ concentration was increased, measured by Dynamic Light Scattering.*

| **Analysis of Variance: $TiO_2$ at 48-hour** | | | | | |
|---|---|---|---|---|---|
| **Source** | **DF** | **Sum of Squares** | **Mean Square** | **F Ratio** | **Prob > F** |
| TiO2 | 6 | 448942.1 | 74823.7 | 16.4337 | <.0001* |
| Error | 133 | 605556.9 | 4553.1 | | |
| C. Total | 139 | 1054499 | | | |

| **Means for One-way ANOVA** | | | | | |
|---|---|---|---|---|---|
| **Level** | **Number** | **Mean** | **Std Error** | **Lower 95%** | **Upper 95%** |
| 0.5 | 20 | 109.05 | 15.088 | 79.21 | 138.89 |
| 1 | 20 | 108.1 | 15.088 | 78.26 | 137.94 |
| 3.5 | 20 | 106.4 | 15.088 | 76.56 | 136.24 |
| 5 | 20 | 148.7 | 15.088 | 118.86 | 178.54 |
| 7 | 20 | 183.45 | 15.088 | 153.61 | 213.29 |
| 10.5 | 20 | 245.45 | 15.088 | 215.61 | 275.29 |
| 14 | 20 | 239.8 | 15.088 | 209.96 | 269.64 |

| **Tukey Kramer HSD: $TiO_2$ at 48-hour** | | | |
|---|---|---|---|
| **q*** | **Alpha** | | |
| 2.99398 | 0.05 | | |
| **Connecting Letters Report** | | | |
| **Level** | | **Mean** | **Std Error** |
| 14 | A | 239.8 | 15.088 |
| 10.5 | A | 245.45 | 15.088 |
| 7 | AB | 183.45 | 15.088 |
| 5 | BC | 148.7 | 15.088 |
| 3.5 | C | 106.4 | 15.088 |
| 1 | C | 108.1 | 15.088 |
| 0.5 | C | 109.05 | 15.088 |

*Levels not connected by the same letter are significantly different
*Std Error used a pooled estimate for error variance

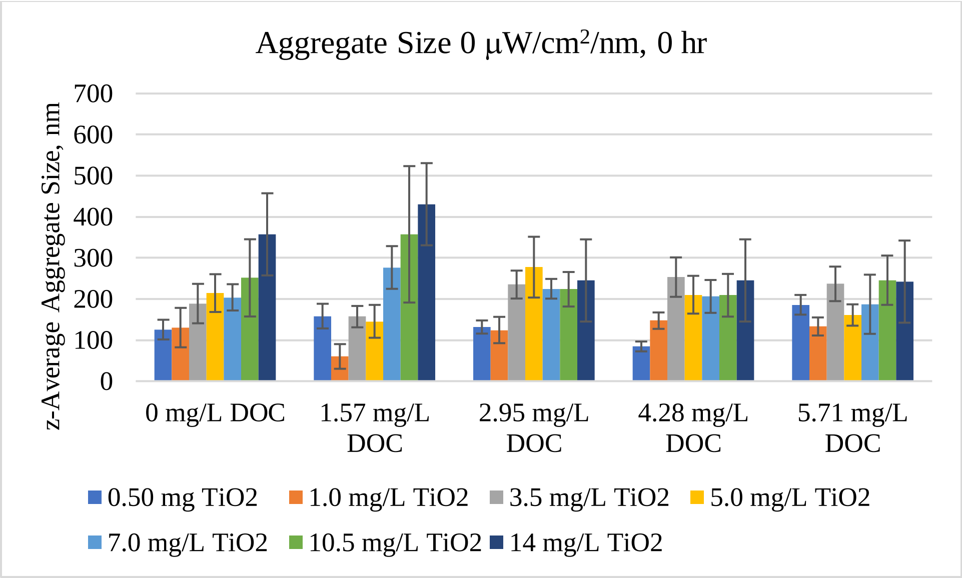


Figure S8. Z-average $TiO_2$ aggregate size under 0 μW/cm2/nm at 0-hour. Any significant differences between treatments are not shown on this graph.

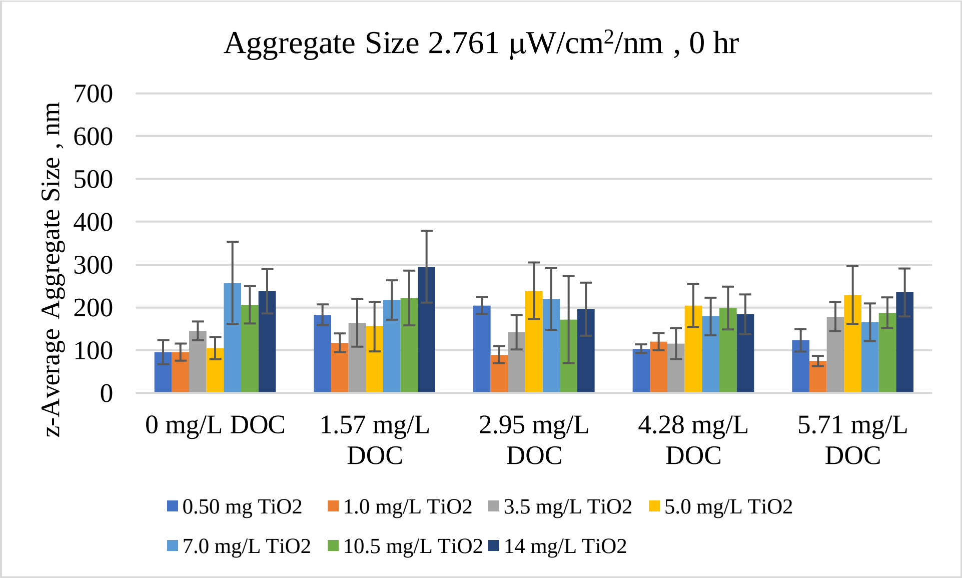


Figure S9. Z-average $TiO_2$ aggregate size under 2.671 μW/cm2/nm at 0-hour. Any significant differences between treatments are not shown on this graph.

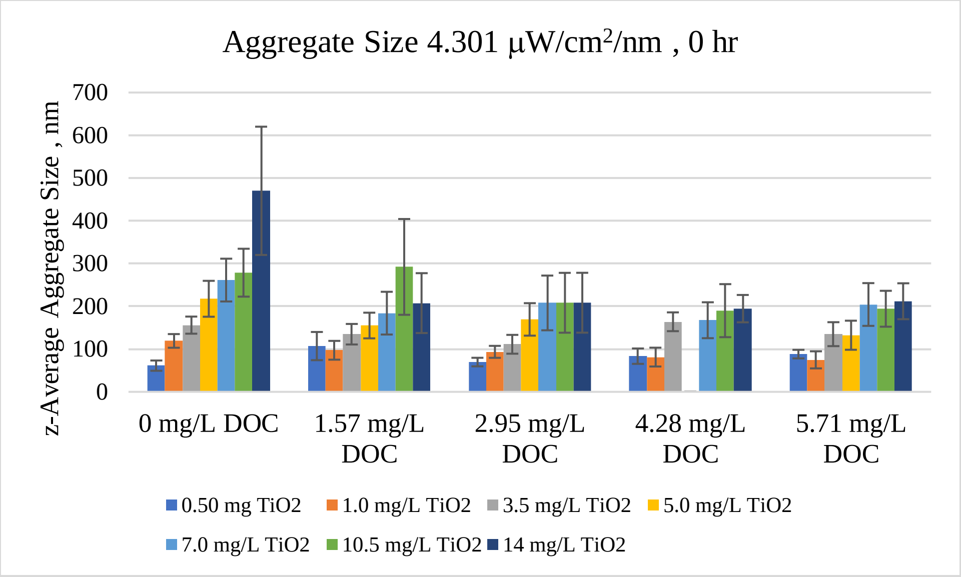


Figure S10. Z-average $TiO_2$ aggregate size under 4.301 μW/cm2/nm at 0-hour. Any significant differences between treatments are not shown on this graph.

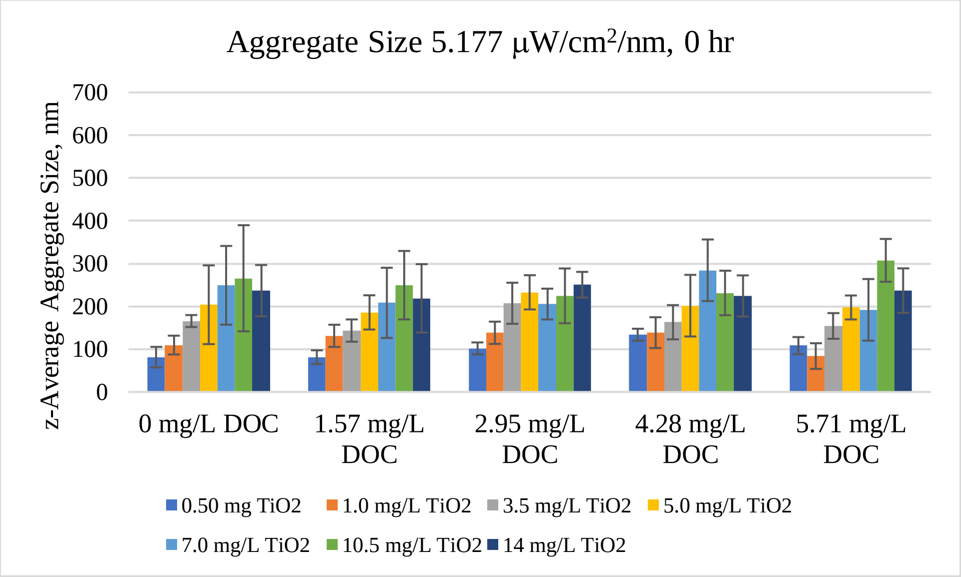


Figure S11. Z-average $TiO_2$ aggregate size under 5.177 μW/cm2/nm at 0-hour. Any significant differences between treatments are not shown on this graph.

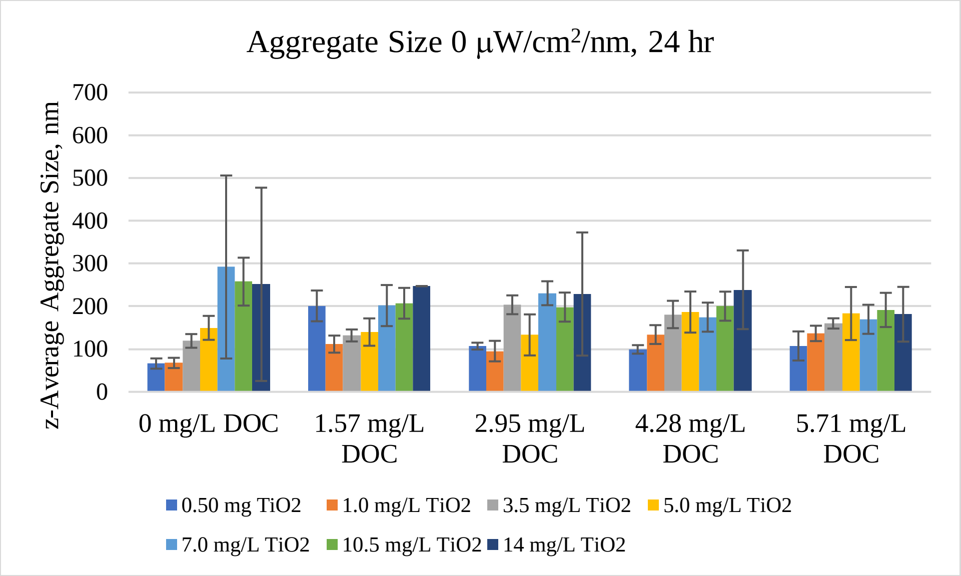


Figure S12. Z-average $TiO_2$ aggregate size under 0 µW/cm2/nm at 24-hour. Any significant differences between treatments are not shown on this graph.

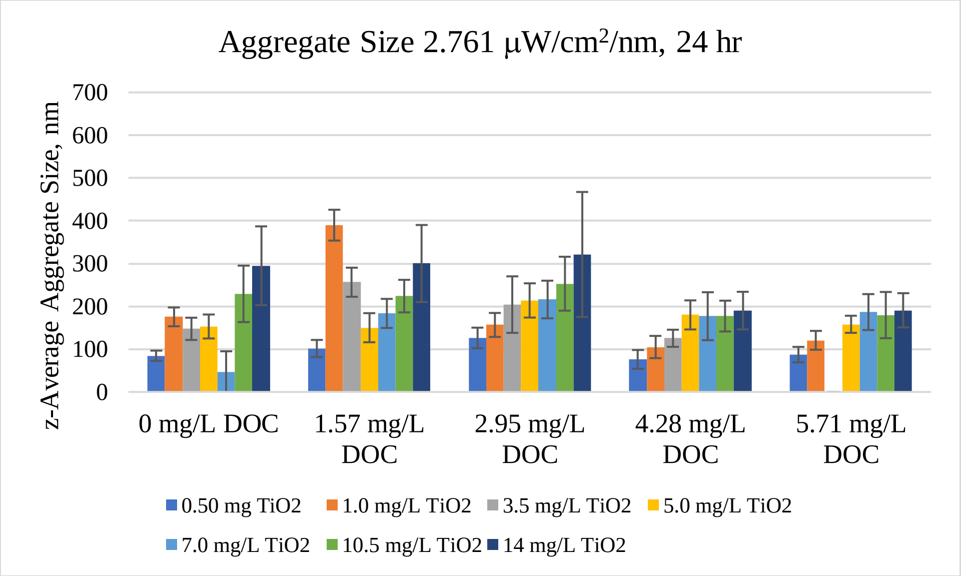


Figure S13. Z-average $TiO_2$ aggregate size under 2.671 µW/cm2/nm at 24-hour. Any significant differences between treatments are not shown on this graph.

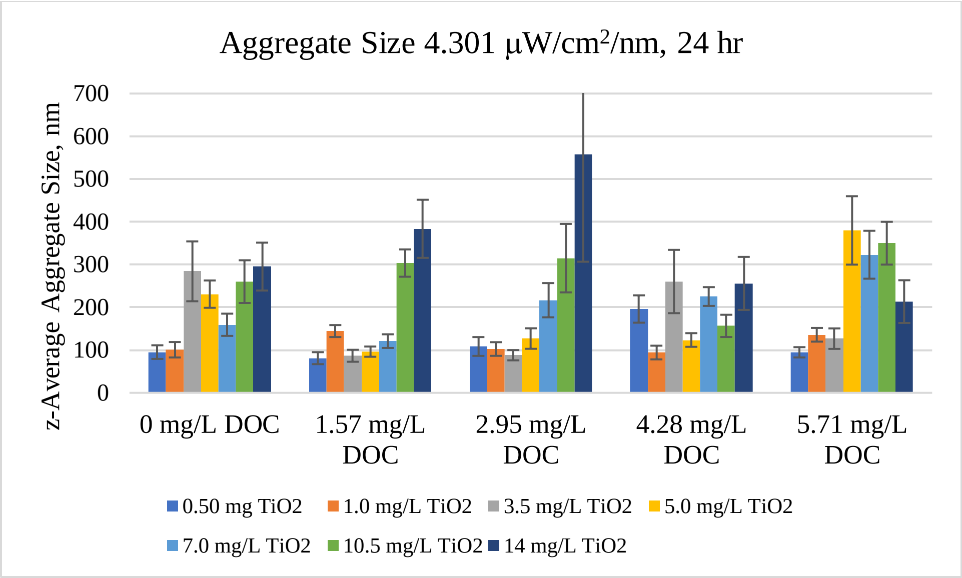


Figure S14. Z-average $TiO_2$ aggregate size under 4.301 μW/cm2/nm at 24-hour. Any significant differences between treatments are not shown on this graph.

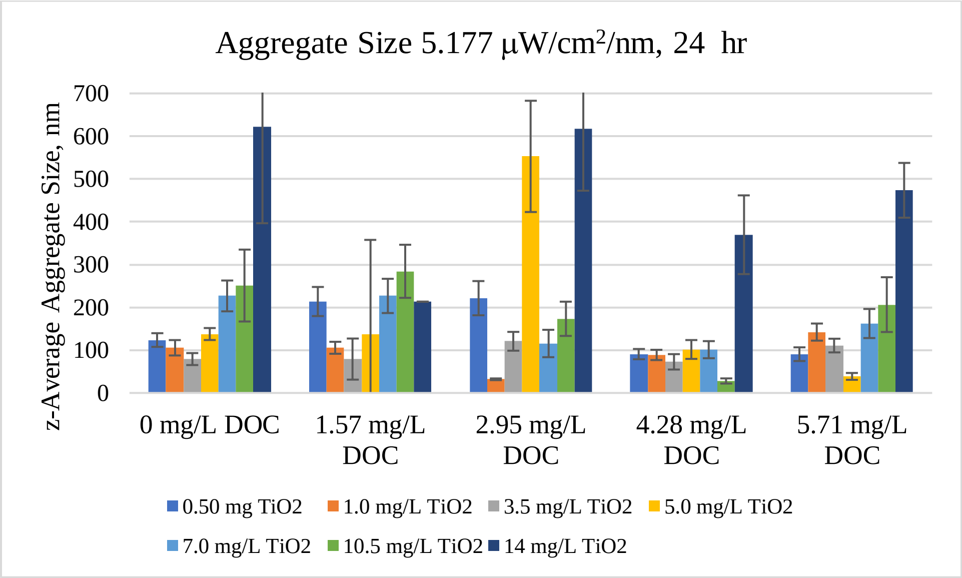


Figure S15. Z-average $TiO_2$ aggregate size under 5.177 μW/cm2/nm at 24-hour. Any significant differences between treatments are not shown on this graph.

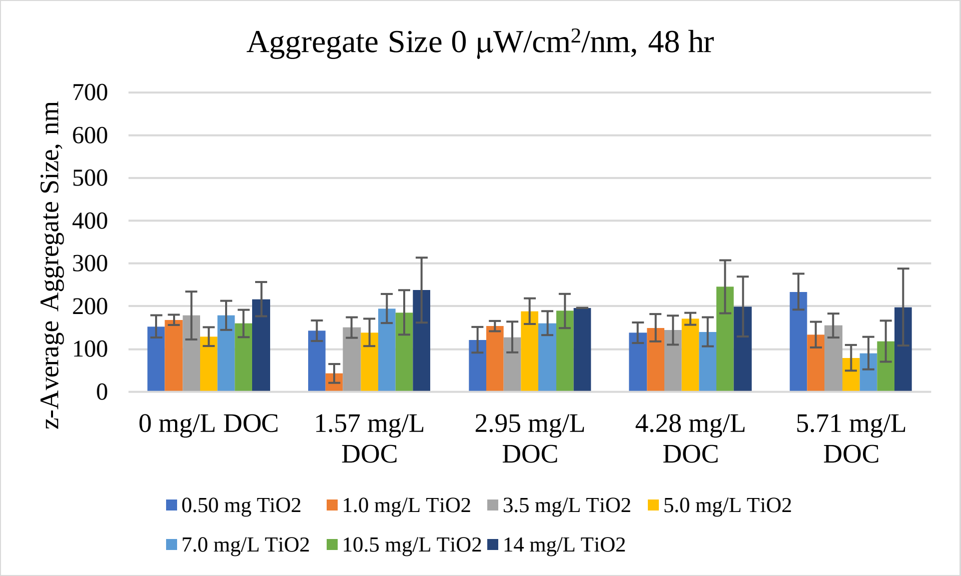


Figure S16. Z-average $TiO_2$ aggregate size under 0 μW/cm2/nm at 48-hour. Any significant differences between treatments are not shown on this graph.

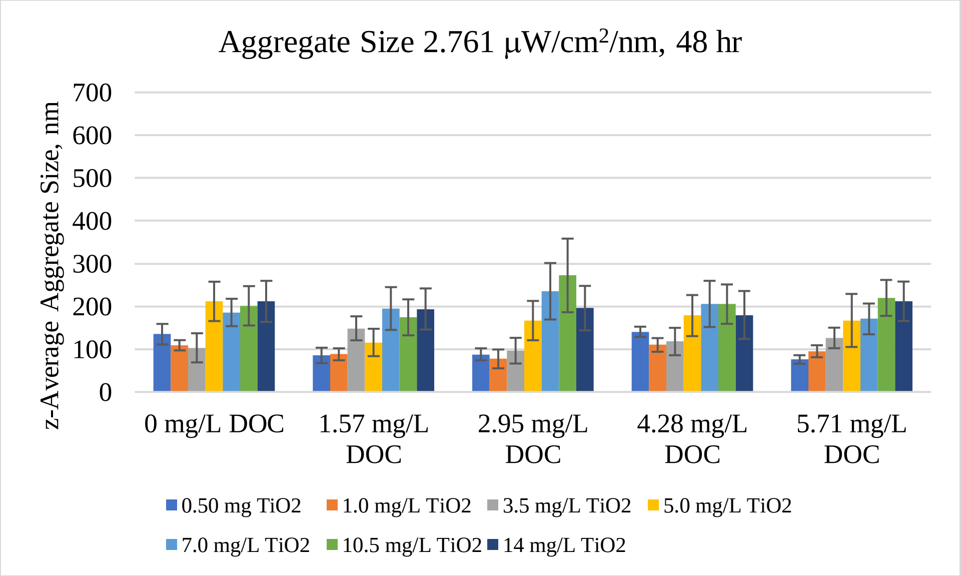


Figure S17. Z-average $TiO_2$ aggregate size under 2.671 μW/cm2/nm at 48-hour. Any significant differences between treatments are not shown on this graph.

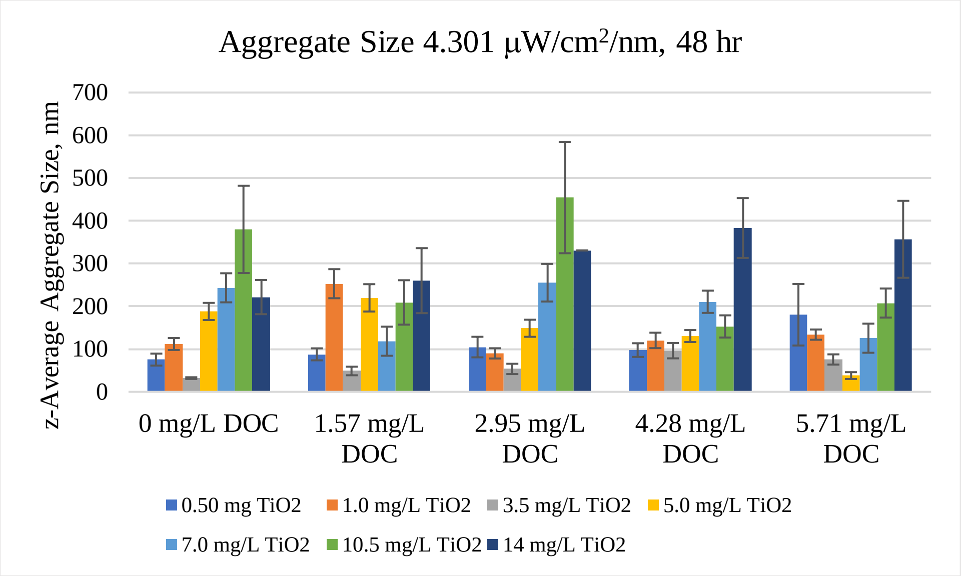


Figure S18. Z-average $TiO_2$ aggregate size under 4.301 μW/$cm^2$/nm at 48-hour. Any significant differences between treatments are not shown on this graph.

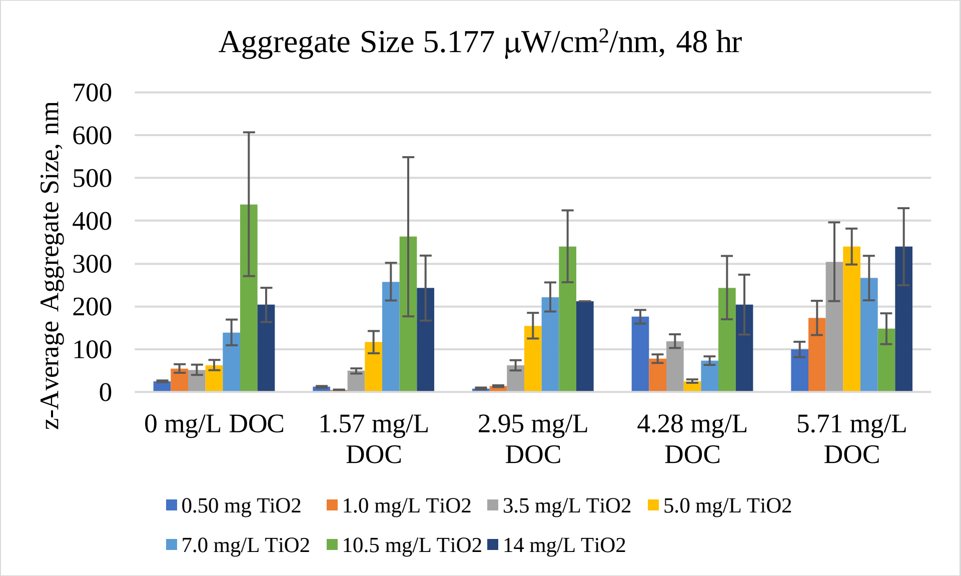


Figure S19. Z-average $TiO_2$ aggregate size under 5.177 μW/$cm^2$/nm at 48-hour. Any significant differences between treatments are not shown on this graph.

Hydroxyl Rate Generation Statistical Summary

Considering the effect that DOC concentration had on hydroxyl radical generation rate per $TiO_2$ at 0 μW/cm2/nm, there were no statistically significant differences at any $TiO_2$ concentration: 0 mg/L: ($F(4,10) = 0$, $p = <1$); 0.5 mg/L: ($F(4,10) = 0.0093$, $p = <.9998$); 1.0 mg/L: ($F(4,10) = 0.2479$, $p = <0.9045$); 3.5 mg/L: ($F(4,10) = 0.0142$, $p = <0.9995$); 5.0 mg/L: ($F(4,10) = 0.008$, $p = <.9998$); 7.0 mg/L: ($F(4,10) = 0.0093$, $p = <.9998$); 10.5 mg/L: ($F(4,10) = 0.1796$, $p = <0.9438$); 14.0 mg/L: ($F(4,10) = 0.7271$, $p = <0.5934$). These results are shown in **Table S28 to S35.**

*Table S28: ANOVA analysis of hydroxyl radical rate generated by 0 mg/L $TiO_2$ under 0 μW/cm2/nm UV-A irradiation in increasing DOC concentration. Measured by fluorescence spectroscopy.*

| **Analysis of Variance: 0 mg/L $TiO_2$ under 0 μW/cm2/nm** | | | | | |
|---|---|---|---|---|---|
| **Source** | **DF** | **Sum of Squares** | **Mean Square** | **F Ratio** | **Prob > F** |
| DOC | 4 | 2.56E-19 | 6.40E-20 | 0 | 1 |
| Error | 10 | 0.00222 | 0.000222 | | |
| C. Total | 14 | 0.00222 | | | |
| **Means for One-way ANOVA** | | | | | |
| **Level** | **Number** | **Mean** | **Std Error** | **Lower 95%** | **Upper 95%** |
| 0 | 3 | -3.33e-10 | 0.0086 | -0.0192 | 0.01917 |
| 1.57 | 3 | -1.81e-20 | 0.0086 | -0.0192 | 0.01917 |
| 2.95 | 3 | 0 | 0.0086 | -0.0192 | 0.01917 |
| 4.28 | 3 | 0 | 0.0086 | -0.0192 | 0.01917 |
| 5.71 | 3 | -3.33e-11 | 0.0086 | -0.0192 | 0.01917 |

*Std Error used a pooled estimate for error variance

*Table S29: ANOVA analysis of hydroxyl radical rate generated by 0.5 mg/L $TiO_2$ under 0 μW/$cm_2$/nm UV-A irradiation in increasing DOC concentration. Measured by fluorescence spectroscopy.*

| **Analysis of Variance: 0.5 mg/L $TiO_2$ under 0 μW/$cm_2$/nm** | | | | | |
|---|---|---|---|---|---|
| **Source** | **DF** | **Sum of Squares** | **Mean Square** | **F Ratio** | **Prob > F** |
| DOC | 4 | 0.000008 | 2.06E-06 | 0.0093 | 0.9998 |
| Error | 10 | 0.002225 | 0.000222 | | |
| C. Total | 14 | 0.002233 | | | |
| **Means for One-way ANOVA** | | | | | |
| **Level** | **Number** | **Mean** | **Std Error** | **Lower 95%** | **Upper 95%** |
| 0 | 3 | 0.0003 | 0.00861 | -0.0189 | 0.0195 |
| 1.57 | 3 | -0.0008 | 0.00861 | -0.0200 | 0.0184 |
| 2.95 | 3 | 0.0001 | 0.00861 | -0.0191 | 0.0193 |
| 4.28 | 3 | -0.0018 | 0.00861 | -0.0210 | 0.0174 |
| 5.71 | 3 | -0.0004 | 0.00861 | -0.0196 | 0.0188 |

*Std Error used a pooled estimate for error variance

*Table S30: ANOVA analysis of hydroxyl radical rate generated by 1.0 mg/L $TiO_2$ under 0 μW/$cm_2$/nm UV-A irradiation in increasing DOC concentration. Measured by fluorescence spectroscopy.*

| **Analysis of Variance: 1.0 mg/L $TiO_2$ under 0 μW/$cm_2$/nm** | | | | | |
|---|---|---|---|---|---|
| **Source** | **DF** | **Sum of Squares** | **Mean Square** | **F Ratio** | **Prob > F** |
| DOC | 4 | 0.00027382 | 0.000068 | 0.2479 | 0.9045 |
| Error | 10 | 0.00276187 | 0.000276 | | |
| C. Total | 14 | 0.00303569 | | | |
| **Means for One-way ANOVA** | | | | | |
| **Level** | **Number** | **Mean** | **Std Error** | **Lower 95%** | **Upper 95%** |
| 0 | 3 | -0.0001 | 0.0096 | -0.0215 | 0.0213 |
| 1.57 | 3 | 0.0099 | 0.0096 | -0.0115 | 0.0312 |
| 2.95 | 3 | 0.0001 | 0.0096 | -0.0213 | 0.0215 |
| 4.28 | 3 | -0.0018 | 0.0096 | -0.0231 | 0.0196 |
| 5.71 | 3 | -0.0011 | 0.0096 | -0.0224 | 0.0203 |

*Std Error used a pooled estimate for error variance

*Table S31: ANOVA analysis of hydroxyl radical rate generated by 3.5 mg/L $TiO_2$ under 0 μW/cm2/nm UV-A irradiation in increasing DOC concentration. Measured by fluorescence spectroscopy.*

| Analysis of Variance: 3.5 mg/L $TiO_2$ under 0 μW/cm2/nm | | | | | |
|---|---|---|---|---|---|
| **Source** | **DF** | **Sum of Squares** | **Mean Square** | **F Ratio** | **Prob > F** |
| DOC | 4 | 0.000013 | 3.20E-06 | 0.0142 | 0.9995 |
| Error | 10 | 0.002258 | 0.000226 | | |
| C. Total | 14 | 0.002271 | | | |
| **Means for One-way ANOVA** | | | | | |
| **Level** | **Number** | **Mean** | **Std Error** | **Lower 95%** | **Upper 95%** |
| 0 | 3 | 0.0000 | 0.0087 | -0.0193 | 0.0193 |
| 1.57 | 3 | 0.0014 | 0.0087 | -0.0179 | 0.0208 |
| 2.95 | 3 | 0.0007 | 0.0087 | -0.0186 | 0.0200 |
| 4.28 | 3 | -0.0014 | 0.0087 | -0.0207 | 0.0180 |
| 5.71 | 3 | 0.0002 | 0.0087 | -0.0191 | 0.0196 |

*Std Error used a pooled estimate for error variance

*Table S32: ANOVA analysis of hydroxyl radical rate generated by 5.0 mg/L $TiO_2$ under 0 μW/cm2/nm UV-A irradiation in increasing DOC concentration. Measured by fluorescence spectroscopy.*

| Analysis of Variance: 5.0 mg/ $TiO_2$ under 0 μW/cm2/nm | | | | | |
|---|---|---|---|---|---|
| **Source** | **DF** | **Sum of Squares** | **Mean Square** | **F Ratio** | **Prob > F** |
| DOC | 4 | 0.000007 | 1.76E-06 | 0.008 | 0.9998 |
| Error | 10 | 0.002191 | 0.000219 | | |
| C. Total | 14 | 0.002198 | | | |
| **Means for One-way ANOVA** | | | | | |
| **Level** | **Number** | **Mean** | **Std Error** | **Lower 95%** | **Upper 95%** |
| 0 | 3 | 0.0011 | 0.0086 | -0.0179 | 0.0202 |
| 1.57 | 3 | 0.0007 | 0.0086 | -0.0183 | 0.0198 |
| 2.95 | 3 | 0.0006 | 0.0086 | -0.0185 | 0.0196 |
| 4.28 | 3 | -0.0008 | 0.0086 | -0.0198 | 0.0183 |
| 5.71 | 3 | -0.0003 | 0.0086 | -0.0193 | 0.0188 |

*Std Error used a pooled estimate for error variance

*Table S33: ANOVA analysis of hydroxyl radical rate generated by 7.0 mg/L $TiO_2$ under 0 μW/cm2/nm UV-A irradiation in increasing DOC concentration. Measured by fluorescence spectroscopy.*

| Analysis of Variance: 7.0 mg/ $TiO_2$ under 0 μW/cm2/nm | | | | | |
|---|---|---|---|---|---|
| **Source** | **DF** | **Sum of Squares** | **Mean Square** | **F Ratio** | **Prob > F** |
| DOC | 4 | 0.0000085 | 2.12E-06 | 0.0093 | 0.9998 |
| Error | 10 | 0.0022679 | 0.000227 | | |
| C. Total | 14 | 0.0022763 | | | |
| **Means for One-way ANOVA** | | | | | |
| **Level** | **Number** | **Mean** | **Std Error** | **Lower 95%** | **Upper 95%** |
| 0 | 3 | 0.0022 | 0.0087 | -0.0171 | 0.02161 |
| 1.57 | 3 | 0.0010 | 0.0087 | -0.0183 | 0.0204 |
| 2.95 | 3 | 0.0005 | 0.0087 | -0.0188 | 0.01992 |
| 4.28 | 3 | 0.0004 | 0.0087 | -0.0190 | 0.01974 |
| 5.71 | 3 | 0.0001 | 0.0087 | -0.0193 | 0.01949 |

*Std Error used a pooled estimate for error variance

*Table S34: ANOVA analysis of hydroxyl radical rate generated by 10.5 mg/L $TiO_2$ under 0 μW/cm2/nm UV-A irradiation in increasing DOC concentration. Measured by fluorescence spectroscopy.*

| Analysis of Variance: 10.5 mg/L $TiO_2$ under 0 μW/cm2/nm | | | | | |
|---|---|---|---|---|---|
| **Source** | **DF** | **Sum of Squares** | **Mean Square** | **F Ratio** | **Prob > F** |
| DOC | 4 | 0.000172 | 0.000043 | 0.1796 | 0.9438 |
| Error | 10 | 0.002390 | 0.000239 | | |
| C. Total | 14 | 0.002561 | | | |
| **Means for One-way ANOVA** | | | | | |
| **Level** | **Number** | **Mean** | **Std Error** | **Lower 95%** | **Upper 95%** |
| 0 | 3 | 0.0007 | 0.0089 | -0.0192 | 0.02055 |
| 1.57 | 3 | 0.0017 | 0.0089 | -0.0182 | 0.02159 |
| 2.95 | 3 | 0.0006 | 0.0089 | -0.0193 | 0.02049 |
| 4.28 | 3 | 0.0009 | 0.0089 | -0.0190 | 0.02076 |
| 5.71 | 3 | 0.0094 | 0.0089 | -0.0105 | 0.02925 |

*Std Error used a pooled estimate for error variance

*Table S35: ANOVA analysis of hydroxyl radical rate generated by 14.0 mg/L $TiO_2$ under 0 μW/cm2/nm UV-A irradiation in increasing DOC concentration. Measured by fluorescence spectroscopy.*

| Analysis of Variance: 14.0 mg/L $TiO_2$ under 0 μW/cm2/nm | | | | | |
|---|---|---|---|---|---|
| **Source** | **DF** | **Sum of Squares** | **Mean Square** | **F Ratio** | **Prob > F** |
| DOC | 4 | 0.0006 | 0.0001 | 0.7271 | 0.5934 |
| Error | 10 | 0.0019 | 0.0002 | | |
| C. Total | 14 | 0.0025 | | | |
| **Means for One-way ANOVA** | | | | | |
| **Level** | **Number** | **Mean** | **Std Error** | **Lower 95%** | **Upper 95%** |
| 0 | 3 | 0.0026 | 0.008 | -0.015 | 0.0204 |
| 1.57 | 3 | 0.0012 | 0.008 | -0.017 | 0.0190 |
| 2.95 | 2 | -0.0158 | 0.010 | -0.038 | 0.0060 |
| 4.28 | 4 | 0.0030 | 0.007 | -0.012 | 0.0185 |
| 5.71 | 3 | 0.0001 | 0.008 | -0.018 | 0.0179 |

*Std Error used a pooled estimate for error variance

Considering the effect that DOC concentration had on hydroxyl radical generation rate per $TiO_2$ at 2.671 μW/cm2/nm, there were statistically significant differences at all $TiO_2$ concentration (except 0 mg/L: ($F(4,10) = 0$, $p$ = <1): 0.5 mg/L: ($F(4,10) = 22.7399$, $p$ = <.0001); 1.0 mg/L: ($F(4,10) = 17.4831$, $p$ = <0.0002); 3.5 mg/L: ($F(4,10) = 95.5329$, $p$ = <0.0001); 5.0 mg/L: ($F(4,10) = 373.8323$, $p$ = <.0001); 7.0 mg/L: ($F(4,10) = 22.747$, $p$ = <.0001); 10.5 mg/L: ($F(4,10) = 28.3497$, $p$ = <0.0001); 14.0 mg/L: ($F(4,10) = 14.879$, $p$ = <0.0003). A *post hoc*

Tukey-Kramer HSD was run to determine levels of significance between the concentrations of DOC per each $TiO_2$ concentration. These results are shown in **Table S36 to S43.**

*Table S36: ANOVA analysis of hydroxyl radical rate generated by 0 mg/L $TiO_2$ under 2.671 μW/$cm_2$/nm UV-A irradiation in increasing DOC concentration. Measured by fluorescence spectroscopy.*

| **Analysis of Variance: 0 mg/L $TiO_2$ under 2.671 μW/$cm_2$/nm** | | | | | |
|---|---|---|---|---|---|
| **Source** | **DF** | **Sum of Squares** | **Mean Square** | **F Ratio** | **Prob > F** |
| DOC | 4 | 1.71E-19 | 4.27E-20 | 0 | 1 |
| Error | 10 | 9.71E-04 | 0.000097 | | |
| C. Total | 14 | 9.71E-04 | | | |
| **Means for One-way ANOVA** | | | | | |
| **Level** | **Number** | **Mean** | **Std Error** | **Lower 95%** | **Upper 95%** |
| 0 | 3 | 0 | 0.0057 | -0.0127 | 0.0127 |
| 1.57 | 3 | 0 | 0.0057 | -0.0127 | 0.0127 |
| 2.95 | 3 | 0 | 0.0057 | -0.0127 | 0.0127 |
| 4.28 | 3 | 3.6E-20 | 0.0057 | -0.0127 | 0.0127 |
| 5.71 | 3 | -2.7e-10 | 0.0057 | -0.0127 | 0.0127 |

*Levels not connected by the same letter are significantly different
*Std Error used a pooled estimate for error variance

*Table S37: ANOVA analysis and Tukey-Kramer HSD* post hoc *of hydroxyl radical rate generated by 0.5 mg/L $TiO_2$ under 2.671 μW/$cm_2$/nm UV-A irradiation in increasing DOC concentration. Measured by fluorescence spectroscopy.*

| **Analysis of Variance: 0.5 mg/L $TiO_2$ under 2.671 μW/$cm_2$/nm** | | | | | |
|---|---|---|---|---|---|
| **Source** | **DF** | **Sum of Squares** | **Mean Square** | **F Ratio** | **Prob > F** |
| DOC | 4 | 0.00474 | 0.00119 | 22.7399 | <.0001* |
| Error | 10 | 0.00052 | 0.00005 | | |
| C. Total | 14 | 0.00526 | | | |
| **Means for One-way ANOVA** | | | | | |
| **Level** | **Number** | **Mean** | **Std Error** | **Lower 95%** | **Upper 95%** |
| 0 | 3 | 0.0358 | 0.0042 | 0.0265 | 0.0451 |
| 1.57 | 3 | 0.0306 | 0.0042 | 0.0213 | 0.0398 |
| 2.95 | 3 | 0.0051 | 0.0042 | -0.0042 | 0.0144 |
| 4.28 | 3 | -0.0107 | 0.0042 | -0.0200 | -0.0014 |
| 5.71 | 3 | 0.0017 | 0.0042 | -0.0075 | 0.011 |
| **Tukey-Kramer HSD for 0.5 mg/L $TiO_2$ under 2.671 μW/$cm_2$/nm** | | | | | |
| **q*** | **Alpha** | **Level** | | **Mean** | **Std Error** |
| 3.29108 | 0.05 | 0 | A | 0.0358 | 0.0042 |
| | | 1.57 | A | 0.0306 | 0.0042 |
| | | 2.95 | B | 0.0051 | 0.0042 |
| | | 5.71 | B | 0.0017 | 0.0042 |
| | | 4.28 | B | -0.0107 | 0.0042 |

*Levels not connected by the same letter are significantly different
*Std Error used a pooled estimate for error variance

*Table S38: ANOVA analysis and Tukey-Kramer HSD* post hoc *of hydroxyl radical rate generated by 01.0 mg/L $TiO_2$ under 2.671 µW/cm2/nm UV-A irradiation in increasing DOC concentration. Measured by fluorescence spectroscopy.*

| **Analysis of Variance: 1.0 mg/L $TiO_2$ under 2.671 µW/cm2/nm** | | | | | |
|---|---|---|---|---|---|
| **Source** | **DF** | **Sum of Squares** | **Mean Square** | **F Ratio** | **Prob > F** |
| DOC | 4 | 0.00833 | 0.00208 | 17.4831 | 0.0002* |
| Error | 10 | 0.00119 | 0.00012 | | |
| C. Total | 14 | 0.00952 | | | |
| **Means for One-way ANOVA** | | | | | |
| **Level** | **Number** | **Mean** | **Std Error** | **Lower 95%** | **Upper 95%** |
| 0 | 3 | 0.0636 | 0.0063 | 0.0496 | 0.07768 |
| 1.57 | 3 | 0.0458 | 0.0063 | 0.0317 | 0.05979 |
| 2.95 | 3 | 0.0159 | 0.0063 | 0.0019 | 0.02995 |
| 4.28 | 3 | 0.0064 | 0.0063 | -0.0076 | 0.02045 |
| 5.71 | 3 | 0.0038 | 0.0063 | -0.0102 | 0.01787 |
| **Tukey-Kramer HSD for 1.0 mg/L $TiO_2$ under 2.671 µW/cm2/nm** | | | | | |
| **q*** | **Alpha** | **Level** | | **Mean** | **Std Error** |
| 3.29108 | 0.05 | 0 | A | 0.0636 | 0.0063 |
| | | 1.57 | A | 0.0457 | 0.0063 |
| | | 2.95 | B | 0.0159 | 0.0063 |
| | | 4.28 | B | 0.0064 | 0.0063 |
| | | 5.71 | B | 0.0038 | 0.0063 |

*Levels not connected by the same letter are significantly different
*Std Error used a pooled estimate for error variance

*Table S39: ANOVA analysis and Tukey-Kramer HSD* post hoc *of hydroxyl radical rate generated by 3.5 mg/L $TiO_2$ under 2.671 μW/cm2/nm UV-A irradiation in increasing DOC concentration. Measured by fluorescence spectroscopy.*

| **Analysis of Variance: 3.5 mg/L $TiO_2$ under 2.671 μW/cm2/nm** | | | | | |
|---|---|---|---|---|---|
| **Source** | **DF** | **Sum of Squares** | **Mean Square** | **F Ratio** | **Prob > F** |
| DOC | 4 | 0.017367 | 0.00434 | 92.5329 | <.0001* |
| Error | 10 | 0.000469 | 0.00005 | | |
| C. Total | 14 | 0.017836 | | | |
| **Means for One-way ANOVA** | | | | | |
| **Level** | **Number** | **Mean** | **Std Error** | **Lower 95%** | **Upper 95%** |
| 0 | 3 | 0.1080 | 0.0040 | 0.0992 | 0.1168 |
| 1.57 | 3 | 0.0956 | 0.0040 | 0.0867 | 0.1044 |
| 2.95 | 3 | 0.0772 | 0.0040 | 0.0684 | 0.0860 |
| 4.28 | 3 | 0.0383 | 0.0040 | 0.0295 | 0.0471 |
| 5.71 | 3 | 0.0183 | 0.0040 | 0.0095 | 0.0271 |
| **Tukey-Kramer HSD for 3.5 mg/L $TiO_2$ under 2.671 μW/cm2/nm** | | | | | |
| **q*** | **Alpha** | **Level** | | **Mean** | **Std Error** |
| 3.29108 | 0.05 | 0 | A | 0.1080 | 0.0040 |
| | | 1.57 | AB | 0.0956 | 0.0040 |
| | | 2.95 | B | 0.0772 | 0.0040 |
| | | 4.28 | C | 0.0383 | 0.0040 |
| | | 5.71 | D | 0.0183 | 0.0040 |

*Levels not connected by the same letter are significantly different

*Std Error used a pooled estimate for error variance

*Table S40: ANOVA analysis and Tukey-Kramer HSD* post hoc *of hydroxyl radical rate generated by 5.0 mg/L $TiO_2$ under 2.671 µW/cm2/nm UV-A irradiation in increasing DOC concentration. Measured by fluorescence spectroscopy.*

| **Analysis of Variance: 5.0 mg/L $TiO_2$ under 2.671 µW/cm2/nm** | | | | | |
|---|---|---|---|---|---|
| **Source** | **DF** | **Sum of Squares** | **Mean Square** | **F Ratio** | **Prob > F** |
| DOC | 4 | 0.023908 | 0.005977 | 373.8323 | <.0001* |
| Error | 10 | 0.000160 | 0.000016 | | |
| C. Total | 14 | 0.024068 | | | |
| **Means for One-way ANOVA** | | | | | |
| **Level** | **Number** | **Mean** | **Std Error** | **Lower 95%** | **Upper 95%** |
| 0 | 3 | 0.1109 | 0.0023 | 0.1058 | 0.1161 |
| 1.57 | 3 | 0.1255 | 0.0023 | 0.1204 | 0.1307 |
| 2.95 | 3 | 0.1234 | 0.0023 | 0.1183 | 0.1286 |
| 4.28 | 3 | 0.0549 | 0.0023 | 0.0498 | 0.0601 |
| 5.71 | 3 | 0.0273 | 0.0023 | 0.0221 | 0.0324 |
| **Tukey-Kramer HSD for 5.0 mg/L $TiO_2$ under 2.671 µW/cm2/nm** | | | | | |
| **q*** | **Alpha** | **Level** | | **Mean** | **Std Error** |
| 3.29108 | 0.05 | 0 | A | 0.1109 | 0.0023 |
| | | 1.57 | B | 0.1255 | 0.0023 |
| | | 2.95 | B | 0.1234 | 0.0023 |
| | | 4.28 | C | 0.0549 | 0.0023 |
| | | 5.71 | D | 0.0273 | 0.0023 |

*Levels not connected by the same letter are significantly different
*Std Error used a pooled estimate for error variance

*Table S41: ANOVA analysis and Tukey-Kramer HSD* post hoc *of hydroxyl radical rate generated by 7.0 mg/L $TiO_2$ under 2.671 µW/cm2/nm UV-A irradiation in increasing DOC concentration. Measured by fluorescence spectroscopy.*

| Analysis of Variance: 7.0 mg/ $TiO_2$ under 2.671 µW/cm2/nm | | | | | |
|---|---|---|---|---|---|
| **Source** | **DF** | **Sum of Squares** | **Mean Square** | **F Ratio** | **Prob > F** |
| DOC | 4 | 0.02893 | 0.00723 | 22.747 | <.0001* |
| Error | 10 | 0.00318 | 0.00032 | | |
| C. Total | 14 | 0.03211 | | | |
| **Means for One-way ANOVA** | | | | | |
| **Level** | **Number** | **Mean** | **Std Error** | **Lower 95%** | **Upper 95%** |
| 0 | 3 | 0.1974 | 0.0103 | 0.1745 | 0.2203 |
| 1.57 | 3 | 0.1884 | 0.0103 | 0.1655 | 0.2114 |
| 2.95 | 3 | 0.1821 | 0.0103 | 0.1592 | 0.2050 |
| 4.28 | 3 | 0.1300 | 0.0103 | 0.1070 | 0.1529 |
| 5.71 | 3 | 0.0817 | 0.0103 | 0.0588 | 0.1046 |
| **Tukey-Kramer HSD for 7.0 mg/L $TiO_2$ under 2.671 µW/cm2/nm** | | | | | |
| **q*** | **Alpha** | **Level** | | **Mean** | **Std Error** |
| 3.29108 | 0.05 | 0 | A | 0.1974 | 0.0103 |
| | | 1.57 | A | 0.1884 | 0.0103 |
| | | 2.95 | A | 0.1821 | 0.0103 |
| | | 4.28 | B | 0.1300 | 0.0103 |
| | | 5.71 | C | 0.0817 | 0.0103 |

*Levels not connected by the same letter are significantly different
*Std Error used a pooled estimate for error variance

*Table S42: ANOVA analysis and Tukey-Kramer HSD* post hoc *of hydroxyl radical rate generated by 10.5 mg/L $TiO_2$ under 2.671 μW/cm2/nm UV-A irradiation in increasing DOC concentration. Measured by fluorescence spectroscopy.*

| **Analysis of Variance: 10.5 mg/L $TiO_2$ under 2.671 μW/cm2/nm** | | | | | |
|---|---|---|---|---|---|
| **Source** | **DF** | **Sum of Squares** | **Mean Square** | **F Ratio** | **Prob > F** |
| DOC | 4 | 0.04948 | 0.01237 | 28.3497 | <.0001* |
| Error | 10 | 0.00436 | 0.00044 | | |
| C. Total | 14 | 0.05385 | | | |
| **Means for One-way ANOVA** | | | | | |
| **Level** | **Number** | **Mean** | **Std Error** | **Lower 95%** | **Upper 95%** |
| 0 | 3 | 0.3055 | 0.0121 | 0.2787 | 0.3324 |
| 1.57 | 3 | 0.3095 | 0.0121 | 0.2827 | 0.3364 |
| 2.95 | 3 | 0.2689 | 0.0121 | 0.2420 | 0.2958 |
| 4.28 | 3 | 0.2301 | 0.0121 | 0.2033 | 0.2570 |
| 5.71 | 3 | 0.1542 | 0.0121 | 0.1273 | 0.1810 |
| **Tukey-Kramer HSD for 10.5 mg/L $TiO_2$ under 2.671 μW/cm2/nm** | | | | | |
| **q*** | **Alpha** | **Level** | | **Mean** | **Std Error** |
| 3.29108 | 0.05 | 0 | A | 0.3055237 | 0.01206 |
| | | 1.57 | A | 0.30952804 | 0.01206 |
| | | 2.95 | AB | 0.26887861 | 0.01206 |
| | | 4.28 | B | 0.23013971 | 0.01206 |
| | | 5.71 | C | 0.15416975 | 0.01206 |

*Levels not connected by the same letter are significantly different
*Std Error used a pooled estimate for error variance

*Table S43: ANOVA analysis and Tukey-Kramer HSD* post hoc *of hydroxyl radical rate generated by 14.0 mg/L $TiO_2$ under 2.671 μW/cm2/nm UV-A irradiation in increasing DOC concentration. Measured by fluorescence spectroscopy.*

| **Analysis of Variance: 14.0 mg/L $TiO_2$ under 2.671 μW/cm2/nm** | | | | | |
|---|---|---|---|---|---|
| **Source** | **DF** | **Sum of Squares** | **Mean Square** | **F Ratio** | **Prob > F** |
| DOC | 4 | 0.030287 | 0.00757 | 14.879 | 0.0003* |
| Error | 10 | 0.005089 | 0.00051 | | |
| C. Total | 14 | 0.035376 | | | |
| **Means for One-way ANOVA** | | | | | |
| **Level** | **Number** | **Mean** | **Std Error** | **Lower 95%** | **Upper 95%** |
| 0 | 3 | 0.4022 | 0.0130 | 0.3731 | 0.4312 |
| 1.57 | 3 | 0.4009 | 0.0130 | 0.3719 | 0.4299 |
| 2.95 | 3 | 0.3880 | 0.0130 | 0.3589 | 0.4170 |
| 4.28 | 3 | 0.3525 | 0.0130 | 0.3235 | 0.3815 |
| 5.71 | 3 | 0.2829 | 0.0130 | 0.2539 | 0.3119 |
| **Tukey-Kramer HSD for 14.0 mg/L $TiO_2$ under 2.671 μW/cm2/nm** | | | | | |
| **q*** | **Alpha** | **Level** | | **Mean** | **Std Error** |
| 3.29108 | 0.05 | 0 | A | 0.4022 | 0.0130 |
| | | 1.57 | A | 0.4009 | 0.0130 |
| | | 2.95 | A | 0.3880 | 0.0130 |
| | | 4.28 | A | 0.3525 | 0.0130 |
| | | 5.71 | B | 0.2829 | 0.0130 |

*Levels not connected by the same letter are significantly different
*Std Error used a pooled estimate for error variance

Considering the effect that DOC concentration had on hydroxyl radical generation rate per $TiO_2$ at 4.301 μW/cm2/nm, there were statistically significant differences at all $TiO_2$ concentration (except 0 mg/L: ($F(4,10) = 0$, $p = <1$):
0.5 mg/L: ($F(4,10) = 115.8457$, $p = <.0001$); 1.0 mg/L: ($F(4,10) = 215.9192$, $p = <0.0001$);
3.5 mg/L: ($F(4,10) = 9.379$, $p = <0.0020$); 5.0 mg/L: ($F(4,10) = 369.9601$, $p = <.0001$);
7.0 mg/L: ($F(4,10) = 8.4766$, $p = <.0030$); 10.5 mg/L: ($F(4,10) = 39.2287$, $p = <0.0001$); 14.0 mg/L: ($F(4,10) = 52.8828$, $p = <0.0003$). A *post hoc* Tukey-Kramer HSD was run to determine levels of significance between the concentrations of DOC per each $TiO_2$ concentration. These results are shown in **Table S44 to S51.**

*Table S44: ANOVA analysis of hydroxyl radical rate generated by 0 mg/L $TiO_2$ under 4.301 μW/cm2/nm UV-A irradiation in increasing DOC concentration. Measured by fluorescence spectroscopy.*

| Analysis of Variance: 0 mg/L $TiO_2$ under 4.301 μW/cm2/nm | | | | | |
|---|---|---|---|---|---|
| **Source** | **DF** | **Sum of Squares** | **Mean Square** | **F Ratio** | **Prob > F** |
| DOC | 4 | 4.27E-20 | 1.07E-20 | 0 | 1 |
| Error | 10 | 0.000037 | 3.76E-06 | | |
| C. Total | 14 | 0.000037 | | | |
| **Means for Oneway Anova** | | | | | |
| **Level** | **Number** | **Mean** | **Std Error** | **Lower 95%** | **Upper 95%** |
| 0 | 3 | -1.45e-19 | 0.00112 | -0.0025 | 0.00249 |
| 1.57 | 3 | -7.23e-20 | 0.00112 | -0.0025 | 0.00249 |
| 2.95 | 3 | 7.23E-20 | 0.00112 | -0.0025 | 0.00249 |
| 4.28 | 3 | 0 | 0.00112 | -0.0025 | 0.00249 |
| 5.71 | 3 | 1.33E-10 | 0.00112 | -0.0025 | 0.00249 |

*Std Error used a pooled estimate for error variance

*Table S45: ANOVA analysis and Tukey-Kramer HSD* post hoc *of hydroxyl radical rate generated by 0.5 mg/L $TiO_2$ under 4.301 μW/cm2/nm UV-A irradiation in increasing DOC concentration. Measured by fluorescence spectroscopy.*

| Analysis of Variance: 0.5 mg/L $TiO_2$ under 4.301 μW/cm2/nm | | | | | |
|---|---|---|---|---|---|
| **Source** | **DF** | **Sum of Squares** | **Mean Square** | **F Ratio** | **Prob > F** |
| DOC | 4 | 0.001615 | 0.000404 | 115.8457 | <.0001* |
| Error | 10 | 0.000035 | 3.49E-06 | | |
| C. Total | 14 | 0.001650 | | | |
| **Means for Oneway Anova** | | | | | |
| **Level** | **Number** | **Mean** | **Std Error** | **Lower 95%** | **Upper 95%** |
| 0 | 3 | 0.0294 | 0.00108 | 0.0270 | 0.0318 |
| 1.57 | 3 | 0.0066 | 0.00108 | 0.0042 | 0.0090 |
| 2.95 | 3 | 0.0051 | 0.00108 | 0.0027 | 0.0075 |
| 4.28 | 3 | 0.0022 | 0.00108 | -0.0002 | 0.0046 |
| 5.71 | 3 | 0.0017 | 0.00108 | -0.0007 | 0.0042 |
| **Tukey-Kramer HSD for $TiO_2$ under 4.301 μW/cm2/nm** | | | | | |
| **q*** | **Alpha** | **Level** | | **Mean** | **Std Error** |
| 3.29108 | 0.05 | 0 | A | 0.0294 | 0.00108 |
| | | 1.57 | B | 0.0066 | 0.00108 |
| | | 2.95 | B | 0.0051 | 0.00108 |
| | | 4.28 | B | 0.0022 | 0.00108 |
| | | 5.71 | B | 0.0017 | 0.00108 |

*Levels not connected by the same letter are significantly different
*Std Error used a pooled estimate for error variance

*Table S46: ANOVA analysis and Tukey-Kramer HSD* post hoc *of hydroxyl radical rate generated by 1.0 mg/L $TiO_2$ under 4.301 μW/cm2/nm UV-A irradiation in increasing DOC concentration. Measured by fluorescence spectroscopy.*

| **Analysis of Variance: 1.0 mg/L $TiO_2$ under 4.301 μW/cm2/nm** | | | | | |
|---|---|---|---|---|---|
| **Source** | **DF** | **Sum of Squares** | **Mean Square** | **F Ratio** | **Prob > F** |
| DOC | 4 | 0.005444 | 0.001361 | 215.9192 | <.0001* |
| Error | 10 | 0.000063 | 6.30E-06 | | |
| C. Total | 14 | 0.005507 | | | |
| **Means for Oneway Anova** | | | | | |
| **Level** | **Number** | **Mean** | **Std Error** | **Lower 95%** | **Upper 95%** |
| 0 | 3 | 0.0574 | 0.00145 | 0.0542 | 0.0606 |
| 1.57 | 3 | 0.0201 | 0.00145 | 0.0168 | 0.0233 |
| 2.95 | 3 | 0.0159 | 0.00145 | 0.0127 | 0.0191 |
| 4.28 | 3 | 0.0078 | 0.00145 | 0.0046 | 0.0110 |
| 5.71 | 3 | 0.0041 | 0.00145 | 0.0008 | 0.0073 |
| **Tukey-Kramer HSD for 1.0 mg/L $TiO_2$ under 4.301 μW/cm2/nm** | | | | | |
| **q*** | **Alpha** | **Level** | | **Mean** | **Std Error** |
| 3.29108 | 0.05 | 0 | A | 0.0574 | 0.00145 |
| | | 1.57 | B | 0.0201 | 0.00145 |
| | | 2.95 | B | 0.0159 | 0.00145 |
| | | 4.28 | C | 0.0078 | 0.00145 |
| | | 5.71 | C | 0.0041 | 0.00145 |

*Levels not connected by the same letter are significantly different

*Std Error used a pooled estimate for error variance

*Table S47: ANOVA analysis and Tukey-Kramer HSD* post hoc *of hydroxyl radical rate generated by 3.5 mg/L $TiO_2$ under 4.301 μW/$cm_2$/nm UV-A irradiation in increasing DOC concentration. Measured by fluorescence spectroscopy.*

| **Analysis of Variance: 3.5 mg/L $TiO_2$ under 4.301 μW/$cm_2$/nm** | | | | | |
|---|---|---|---|---|---|
| **Source** | **DF** | **Sum of Squares** | **Mean Square** | **F Ratio** | **Prob > F** |
| DOC | 4 | 0.028179 | 0.007045 | 9.379 | 0.0020* |
| Error | 10 | 0.007511 | 0.000751 | | |
| C. Total | 14 | 0.035690 | | | |
| **Means for Oneway Anova** | | | | | |
| **Level** | **Number** | **Mean** | **Std Error** | **Lower 95%** | **Upper 95%** |
| 0 | 3 | 0.1606 | 0.01582 | 0.12536 | 0.19587 |
| 1.57 | 3 | 0.0876 | 0.01582 | 0.0523 | 0.12282 |
| 2.95 | 3 | X | x | x | x |
| 4.28 | 3 | 0.1095 | 0.01582 | 0.07424 | 0.14475 |
| 5.71 | 3 | 0.0353 | 0.01582 | 3.15E-05 | 0.07054 |
| **Tukey-Kramer HSD for 3.5 mg/L $TiO_2$ under 4.301 μW/$cm_2$/nm** | | | | | |
| **q*** | **Alpha** | **Level** | | **Mean** | **Std Error** |
| 3.29108 | 0.05 | 0 | A | 0.1606 | 0.01582 |
| | | 1.57 | AB | 0.0876 | 0.01582 |
| | | 2.95 | x | X | x |
| | | 4.28 | ABC | 0.1095 | 0.01582 |
| | | 5.71 | C | 0.0353 | 0.01582 |

*Levels not connected by the same letter are significantly different
*Std Error used a pooled estimate for error variance

*Table S48: ANOVA analysis and Tukey-Kramer HSD* post hoc *of hydroxyl radical rate generated by 5.0 mg/L $TiO_2$ under 4.301 µW/$cm^2$/nm UV-A irradiation in increasing DOC concentration. Measured by fluorescence spectroscopy.*

| **Analysis of Variance: 5.0 mg/L $TiO_2$ under 4.301 µW/$cm^2$/nm** | | | | | |
|---|---|---|---|---|---|
| **Source** | **DF** | **Sum of Squares** | **Mean Square** | **F Ratio** | **Prob > F** |
| DOC | 4 | 0.017523 | 0.004381 | 369.9601 | <.0001* |
| Error | 10 | 0.000118 | 0.000012 | | |
| C. Total | 14 | 0.017640 | | | |
| **Means for Oneway Anova** | | | | | |
| **Level** | **Number** | **Mean** | **Std Error** | **Lower 95%** | **Upper 95%** |
| 0 | 3 | x | x | x | x |
| 1.57 | 3 | 0.1152 | 0.00199 | 0.1108 | 0.1197 |
| 2.95 | 3 | 0.1234 | 0.00199 | 0.1190 | 0.1279 |
| 4.28 | 3 | 0.1003 | 0.00199 | 0.0959 | 0.1047 |
| 5.71 | 3 | 0.0326 | 0.00199 | 0.0282 | 0.0370 |
| **Tukey-Kramer HSD for 5.0 mg/L $TiO_2$ under 4.301 µW/$cm^2$/nm** | | | | | |
| **q*** | **Alpha** | **Level** | | **Mean** | **Std Error** |
| 3.29108 | 0.05 | 0 | x | x | x |
| | | 1.57 | A | 0.1152 | 0.00199 |
| | | 2.95 | A | 0.1234 | 0.00199 |
| | | 4.28 | B | 0.1003 | 0.00199 |
| | | 5.71 | C | 0.0326 | 0.00199 |

*Levels not connected by the same letter are significantly different
*Std Error used a pooled estimate for error variance

*Table S49: ANOVA analysis and Tukey-Kramer HSD* post hoc *of hydroxyl radical rate generated by 7.0 mg/L $TiO_2$ under 4.301 μW/$cm_2$/nm UV-A irradiation in increasing DOC concentration. Measured by fluorescence spectroscopy.*

| **Analysis of Variance: 7.0 mg/L $TiO_2$ under 4.301 μW/$cm_2$/nm** | | | | | |
|---|---|---|---|---|---|
| **Source** | **DF** | **Sum of Squares** | **Mean Square** | **F Ratio** | **Prob > F** |
| DOC | 4 | 0.108442 | 0.02711 | 8.4766 | 0.0030* |
| Error | 10 | 0.031983 | 0.003198 | | |
| C. Total | 14 | 0.140425 | | | |
| **Means for Oneway Anova** | | | | | |
| **Level** | **Number** | **Mean** | **Std Error** | **Lower 95%** | **Upper 95%** |
| 0 | 3 | 0.3553 | 0.03265 | 0.2825 | 0.4280 |
| 1.57 | 3 | 0.2575 | 0.03265 | 0.1847 | 0.3302 |
| 2.95 | 3 | 0.2014 | 0.03265 | 0.1287 | 0.2742 |
| 4.28 | 3 | 0.2011 | 0.03265 | 0.1284 | 0.2739 |
| 5.71 | 3 | 0.0945 | 0.03265 | 0.0217 | 0.1672 |
| **Tukey-Kramer HSD for 7.0 mg/L $TiO_2$ under 4.301 μW/$cm_2$/nm** | | | | | |
| **q*** | **Alpha** | **Level** | | **Mean** | **Std Error** |
| 3.29108 | 0.05 | 0 | A | 0.3553 | 0.03265 |
| | | 1.57 | AB | 0.2575 | 0.03265 |
| | | 2.95 | BC | 0.2014 | 0.03265 |
| | | 4.28 | BC | 0.2011 | 0.03265 |
| | | 5.71 | C | 0.0945 | 0.03265 |

*Levels not connected by the same letter are significantly different
*Std Error used a pooled estimate for error variance

*Table S50: ANOVA analysis and Tukey-Kramer HSD* post hoc *of hydroxyl radical rate generated by 10.5 mg/L $TiO_2$ under 4.301 $\mu W/cm^2/nm$ UV-A irradiation in increasing DOC concentration. Measured by fluorescence spectroscopy.*

| **Analysis of Variance: 10.5 mg/L $TiO_2$ under 4.301 $\mu W/cm^2/nm$** | | | | | |
|---|---|---|---|---|---|
| **Source** | **DF** | **Sum of Squares** | **Mean Square** | **F Ratio** | **Prob > F** |
| DOC | 4 | 0.406377 | 0.101594 | 39.2287 | <.0001* |
| Error | 10 | 0.025898 | 0.00259 | | |
| C. Total | 14 | 0.432275 | | | |

| **Means for Oneway Anova** | | | | | |
|---|---|---|---|---|---|
| **Level** | **Number** | **Mean** | **Std Error** | **Lower 95%** | **Upper 95%** |
| 0 | 3 | 0.6614 | 0.02938 | 0.5960 | 0.7269 |
| 1.57 | 3 | 0.4340 | 0.02938 | 0.3685 | 0.4994 |
| 2.95 | 3 | 0.3629 | 0.02938 | 0.2975 | 0.4284 |
| 4.28 | 3 | 0.3013 | 0.02938 | 0.2358 | 0.3668 |
| 5.71 | 3 | 0.1633 | 0.02938 | 0.0978 | 0.2287 |

| **Tukey-Kramer HSD for 10.5 mg/L $TiO_2$ under 4.301 $\mu W/cm^2/nm$** | | | | | |
|---|---|---|---|---|---|
| **q*** | **Alpha** | **Level** | | **Mean** | **Std Error** |
| 3.29108 | 0.05 | 0 | A | 0.6614 | 0.02938 |
| | | 1.57 | B | 0.4340 | 0.02938 |
| | | 2.95 | B | 0.3629 | 0.02938 |
| | | 4.28 | B | 0.3013 | 0.02938 |
| | | 5.71 | C | 0.1633 | 0.02938 |

*Levels not connected by the same letter are significantly different
*Std Error used a pooled estimate for error variance

*Table S51: ANOVA analysis and Tukey-Kramer HSD* post hoc *of hydroxyl radical rate generated by 14.0 mg/L $TiO_2$ under 4.301 μW/cm2/nm UV-A irradiation in increasing DOC concentration. Measured by fluorescence spectroscopy.*

| Analysis of Variance: 14.0 mg/L $TiO_2$ under 4.301 μW/cm2/nm | | | | | |
|---|---|---|---|---|---|
| **Source** | **DF** | **Sum of Squares** | **Mean Square** | **F Ratio** | **Prob > F** |
| DOC | 4 | 0.520720 | 0.13018 | 52.8828 | <.0001* |
| Error | 10 | 0.024617 | 0.002462 | | |
| C. Total | 14 | 0.545337 | | | |
| **Means for Oneway Anova** | | | | | |
| **Level** | **Number** | **Mean** | **Std Error** | **Lower 95%** | **Upper 95%** |
| 0 | 3 | 0.8179 | 0.02865 | 0.7540 | 0.8817 |
| 1.57 | 3 | 0.6005 | 0.02865 | 0.5367 | 0.6643 |
| 2.95 | 3 | 0.5960 | 0.02865 | 0.5322 | 0.6599 |
| 4.28 | 3 | 0.4237 | 0.02865 | 0.3598 | 0.4875 |
| 5.71 | 3 | 0.2642 | 0.02865 | 0.2004 | 0.3281 |

| Tukey-Kramer HSD for 14.0 mg/L $TiO_2$ under 4.301 μW/cm2/nm | | | | | |
|---|---|---|---|---|---|
| **q*** | **Alpha** | **Level** | | **Mean** | **Std Error** |
| 3.29108 | 0.05 | 0 | A | 0.8179 | 0.02865 |
| | | 1.57 | B | 0.6005 | 0.02865 |
| | | 2.95 | B | 0.5960 | 0.02865 |
| | | 4.28 | C | 0.4237 | 0.02865 |
| | | 5.71 | D | 0.2642 | 0.02865 |

*Levels not connected by the same letter are significantly different
*Std Error used a pooled estimate for error variance

Considering the effect that DOC concentration had on hydroxyl radical generation rate per $TiO_2$ at 5.177 μW/cm2/nm, there were statistically significant differences at all $TiO_2$ concentration (except 0 mg/L: ($F(4,10) = 0$, $p$ = <1): 0.5 mg/L: ($F(4,10) = 18.211$, $p$ = <.0001); 1.0 mg/L: ($F(4,10) = 94.4584$, $p$ = <0.0001);
3.5 mg/L: ($F(4,10) = 41.6108$, $p$ = <0.0001); 5.0 mg/L: ($F(4,10) = 76.1449$, $p$ = <.0001);
7.0 mg/L: ($F(4,10) = 8175.8544$, $p$ = <.0001); 10.5 mg/L: ($F(4,10) = 103.0842$, $p$ = <0.0001);
14.0 mg/L: ($F(4,10) = 30.1188$, $p$ = <0.0001). A *post hoc* Tukey-Kramer HSD was run to determine levels of significance between the concentrations of DOC per each $TiO_2$ concentration. These results are shown in **Table S52 to S.**

*Table S52: ANOVA analysis of hydroxyl radical rate generated by 0 mg/L $TiO_2$ under 5.177 μW/cm2/nm UV-A irradiation in increasing DOC concentration. Measured by fluorescence spectroscopy.*

| **Analysis of Variance: 0 mg/L $TiO_2$ under 5.177 μW/cm2/nm** | | | | | |
|---|---|---|---|---|---|
| **Source** | **DF** | **Sum of Squares** | **Mean Square** | **F Ratio** | **Prob > F** |
| DOC | 4 | 4.00E-19 | 1.00E-19 | 0 | 1 |
| Error | 10 | 4.61E-05 | 4.60E-06 | | |
| C. Total | 14 | 4.61E-05 | | | |
| **Means for One-way ANOVA** | | | | | |
| **Level** | **Number** | **Mean** | **Std Error** | **Lower 95%** | **Upper 95%** |
| 0 | 3 | -7.23e-20 | 0.00124 | -0.0028 | 0.0028 |
| 1.57 | 3 | -3.33e-10 | 0.00124 | -0.0028 | 0.0028 |
| 2.95 | 3 | 3.61E-20 | 0.00124 | -0.0028 | 0.0028 |
| 4.28 | 3 | -3.33e-10 | 0.00124 | -0.0028 | 0.0028 |
| 5.71 | 3 | 0 | 0.00124 | -0.0028 | 0.0028 |

*Std Error used a pooled estimate for error variance

*Table S53: ANOVA analysis and Tukey-Kramer HSD* post hoc *of hydroxyl radical rate generated by 0.5 mg/L $TiO_2$ under 5.177 μW/cm2/nm UV-A irradiation in increasing DOC concentration. Measured by fluorescence spectroscopy.*

| **Analysis of Variance: 0.5 mg/L $TiO_2$ under 5.177 μW/cm2/nm** | | | | | |
|---|---|---|---|---|---|
| **Source** | **DF** | **Sum of Squares** | **Mean Square** | **F Ratio** | **Prob > F** |
| DOC | 4 | 0.001057 | 0.000264 | 18.211 | 0.0001* |
| Error | 10 | 0.000145 | 0.000015 | | |
| C. Total | 14 | 0.001202 | | | |
| **Means for One-way ANOVA** | | | | | |
| **Level** | **Number** | **Mean** | **Std Error** | **Lower 95%** | **Upper 95%** |
| 0 | 3 | 0.0282 | 0.0022 | 0.0233 | 0.0331 |
| 1.57 | 3 | 0.0171 | 0.0022 | 0.0122 | 0.0220 |
| 2.95 | 3 | 0.0084 | 0.0022 | 0.0035 | 0.0133 |
| 4.28 | 3 | 0.0089 | 0.0022 | 0.0040 | 0.0138 |
| 5.71 | 3 | 0.0048 | 0.0022 | -0.0001 | 0.0098 |
| **Tukey-Kramer HSD for 0.5 $TiO_2$ under 5.177 μW/cm2/nm** | | | | | |
| **q*** | **Alpha** | **Level** | | **Mean** | **Std Error** |
| 3.29108 | 0.05 | 0 | A | 0.0282 | 0.0022 |
| | | 1.57 | B | 0.0171 | 0.0022 |
| | | 4.28 | BC | 0.0089 | 0.0022 |
| | | 2.95 | BC | 0.0084 | 0.0022 |
| | | 5.71 | C | 0.0048 | 0.0022 |

*Levels not connected by the same letter are significantly different
*Std Error used a pooled estimate for error variance

*Table S54: ANOVA analysis and Tukey-Kramer HSD* post hoc *of hydroxyl radical rate generated by 1.0 mg/L $TiO_2$ under 5.177 μW/cm$_2$/nm UV-A irradiation in increasing DOC concentration. Measured by fluorescence spectroscopy.*

| **Analysis of Variance: 1.0 mg/L $TiO_2$ under 5.177 μW/cm$_2$/nm** | | | | | |
|---|---|---|---|---|---|
| **Source** | **DF** | **Sum of Squares** | **Mean Square** | **F Ratio** | **Prob > F** |
| DOC | 4 | 0.005793 | 0.001448 | 94.4584 | <.0001* |
| Error | 10 | 0.000153 | 0.000015 | | |
| C. Total | 14 | 0.005947 | | | |
| **Means for One-way ANOVA** | | | | | |
| **Level** | **Number** | **Mean** | **Std Error** | **Lower 95%** | **Upper 95%** |
| 0 | 3 | 0.0609 | 0.0023 | 0.0559 | 0.0660 |
| 1.57 | 3 | 0.0488 | 0.0023 | 0.0438 | 0.0539 |
| 2.95 | 3 | 0.0190 | 0.0023 | 0.0140 | 0.0241 |
| 4.28 | 3 | 0.0179 | 0.0023 | 0.0128 | 0.0229 |
| 5.71 | 3 | 0.0110 | 0.0023 | 0.0059 | 0.0160 |
| **Tukey-Kramer HSD for 1.0 mg/L $TiO_2$ under 5.177 μW/cm$_2$/nm** | | | | | |
| **q*** | **Alpha** | **Level** | | **Mean** | **Std Error** |
| 3.29108 | 0.05 | 0 | A | 0.0609 | 0.0023 |
| | | 1.57 | B | 0.0488 | 0.0023 |
| | | 2.95 | C | 0.0190 | 0.0023 |
| | | 4.28 | C | 0.0179 | 0.0023 |
| | | 5.71 | C | 0.0109 | 0.0023 |

*Levels not connected by the same letter are significantly different
*Std Error used a pooled estimate for error variance

*Table S55: ANOVA analysis and Tukey-Kramer HSD* post hoc *of hydroxyl radical rate generated by 3.5 mg/L $TiO_2$ under 5.177 μW/cm$_2$/nm UV-A irradiation in increasing DOC concentration. Measured by fluorescence spectroscopy.*

| **Analysis of Variance: 3.5 mg/L $TiO_2$ under 5.177 μW/cm2/nm** | | | | | |
|---|---|---|---|---|---|
| **Source** | **DF** | **Sum of Squares** | **Mean Square** | **F Ratio** | **Prob > F** |
| DOC | 4 | 0.376863 | 0.094216 | 41.6108 | <.0001* |
| Error | 10 | 0.022642 | 0.002264 | | |
| C. Total | 14 | 0.399505 | | | |
| **Means for One-way ANOVA** | | | | | |
| **Level** | **Number** | **Mean** | **Std Error** | **Lower 95%** | **Upper 95%** |
| 0 | 3 | 0.4679 | 0.0275 | 0.4066 | 0.5291 |
| 1.57 | 3 | 0.2070 | 0.0275 | 0.1457 | 0.2682 |
| 2.95 | 3 | 0.1097 | 0.0275 | 0.0485 | 0.1709 |
| 4.28 | 3 | 0.0420 | 0.0275 | -0.0192 | 0.1033 |
| 5.71 | 3 | 0.0450 | 0.0275 | -0.0162 | 0.1062 |
| **Tukey-Kramer HSD for 3.5 mg/L $TiO_2$ under 5.177 μW/cm2/nm** | | | | | |
| **q*** | **Alpha** | **Level** | | **Mean** | **Std Error** |
| 3.29108 | 0.05 | 0 | A | 0.4679 | 0.0275 |
| | | 1.57 | B | 0.2070 | 0.0275 |
| | | 2.95 | BC | 0.1097 | 0.0275 |
| | | 4.28 | C | 0.0420 | 0.0275 |
| | | 5.71 | C | 0.0450 | 0.0275 |

*Levels not connected by the same letter are significantly different
*Std Error used a pooled estimate for error variance

*Table S56: ANOVA analysis and Tukey-Kramer HSD* post hoc *of hydroxyl radical rate generated by 5.0 mg/L $TiO_2$ under 5.177 µW/$cm_2$/nm UV-A irradiation in increasing DOC concentration. Measured by fluorescence spectroscopy.*

| **Analysis of Variance: 5.0 mg/L $TiO_2$ under 5.177 µW/$cm_2$/nm** | | | | | |
|---|---|---|---|---|---|
| **Source** | **DF** | **Sum of Squares** | **Mean Square** | **F Ratio** | **Prob > F** |
| DOC | 4 | 0.012385 | 0.003096 | 76.1449 | <.0001* |
| Error | 10 | 0.000407 | 0.000041 | | |
| C. Total | 14 | 0.012791 | | | |
| **Means for One-way ANOVA** | | | | | |
| **Level** | **Number** | **Mean** | **Std Error** | **Lower 95%** | **Upper 95%** |
| 0 | x | x | x | x | x |
| 1.57 | 3 | 0.1473 | 0.0037 | 0.1391 | 0.1555 |
| 2.95 | 3 | 0.1442 | 0.0037 | 0.1360 | 0.1524 |
| 4.28 | 3 | 0.1239 | 0.0037 | 0.1157 | 0.1321 |
| 5.71 | 3 | 0.0674 | 0.0037 | 0.0592 | 0.0756 |
| **Tukey-Kramer HSD for 5.0 mg/L $TiO_2$ under 5.177 µW/$cm_2$/nm** | | | | | |
| **q*** | **Alpha** | **Level** | | **Mean** | **Std Error** |
| 3.29108 | 0.05 | 0 | x | x | x |
| | | 1.57 | A | 0.1473 | 0.0037 |
| | | 2.95 | A | 0.1442 | 0.0037 |
| | | 4.28 | B | 0.1239 | 0.0037 |
| | | 5.71 | C | 0.0674 | 0.0037 |

*Levels not connected by the same letter are significantly different

*Std Error used a pooled estimate for error variance

*Table S57: ANOVA analysis and Tukey-Kramer HSD* post hoc *of hydroxyl radical rate generated by 7.0 mg/L $TiO_2$ under 5.177 μW/cm2/nm UV-A irradiation in increasing DOC concentration. Measured by fluorescence spectroscopy.*

| **Analysis of Variance: 7.0 mg/L $TiO_2$ under 5.177 μW/cm2/nm** | | | | | |
|---|---|---|---|---|---|
| **Source** | **DF** | **Sum of Squares** | **Mean Square** | **F Ratio** | **Prob > F** |
| DOC | 4 | 1.55198 | 0.38799 | 175.8544 | <.0001* |
| Error | 10 | 0.02206 | 0.00221 | | |
| C. Total | 14 | 1.57404 | | | |

| **Means for One-way ANOVA** | | | | | |
|---|---|---|---|---|---|
| **Level** | **Number** | **Mean** | **Std Error** | **Lower 95%** | **Upper 95%** |
| 0 | 3 | 1.0194 | 0.0271 | 0.9590 | 1.0798 |
| 1.57 | 3 | 0.6014 | 0.0271 | 0.5409 | 0.6618 |
| 2.95 | 3 | 0.4814 | 0.0271 | 0.4209 | 0.5418 |
| 4.28 | 3 | 0.2737 | 0.0271 | 0.2132 | 0.3341 |
| 5.71 | 3 | 0.0684 | 0.0271 | 0.0079 | 0.1288 |

| **Tukey-Kramer HSD for 7.0 mg/L $TiO_2$ under 5.177 μW/cm2/nm** | | | | | |
|---|---|---|---|---|---|
| **q*** | **Alpha** | **Level** | | **Mean** | **Std Error** |
| 3.29108 | 0.05 | 0 | A | 1.0194 | 0.0271 |
| | | 1.57 | B | 0.6014 | 0.0271 |
| | | 2.95 | B | 0.4814 | 0.0271 |
| | | 4.28 | C | 0.2737 | 0.0271 |
| | | 5.71 | D | 0.0684 | 0.0271 |

*Levels not connected by the same letter are significantly different
*Std Error used a pooled estimate for error variance

*Table S58: ANOVA analysis and Tukey-Kramer HSD* post hoc *of hydroxyl radical rate generated by 10.5 mg/L $TiO_2$ under 5.177 µW/cm2/nm UV-A irradiation in increasing DOC concentration. Measured by fluorescence spectroscopy.*

| **Analysis of Variance: 10.5 mg/L $TiO_2$ under 5.177 µW/cm2/nm** | | | | | |
|---|---|---|---|---|---|
| **Source** | **DF** | **Sum of Squares** | **Mean Square** | **F Ratio** | **Prob > F** |
| DOC | 4 | 0.761479 | 0.190370 | 103.0842 | <.0001* |
| Error | 10 | 0.018467 | 0.001847 | | |
| C. Total | 14 | 0.779946 | | | |
| **Means for One-way ANOVA** | | | | | |
| **Level** | **Number** | **Mean** | **Std Error** | **Lower 95%** | **Upper 95%** |
| 0 | 3 | 1.1466 | 0.0248 | 1.0913 | 1.2018 |
| 1.57 | 3 | 0.9377 | 0.0248 | 0.8824 | 0.9930 |
| 2.95 | 3 | 0.8395 | 0.0248 | 0.7842 | 0.8947 |
| 4.28 | 3 | 0.5552 | 0.0248 | 0.4999 | 0.6105 |
| 5.71 | 3 | 0.5682 | 0.0248 | 0.5130 | 0.6235 |
| **Tukey-Kramer HSD for 10.5 mg/L $TiO_2$ under 5.177 µW/cm2/nm** | | | | | |
| **q*** | **Alpha** | **Level** | | **Mean** | **Std Error** |
| 3.29108 | 0.05 | 0 | A | 1.1466 | 0.0248 |
| | | 1.57 | B | 0.9377 | 0.0248 |
| | | 2.95 | B | 0.8395 | 0.0248 |
| | | 4.28 | C | 0.5552 | 0.0248 |
| | | 5.71 | C | 0.5682 | 0.0248 |

*Levels not connected by the same letter are significantly different
*Std Error used a pooled estimate for error variance

*Table S59: ANOVA analysis and Tukey-Kramer HSD* post hoc *of hydroxyl radical rate generated by 14.0 mg/L $TiO_2$ under 5.177 $\mu$W/$cm_2$/nm UV-A irradiation in increasing DOC concentration. Measured by fluorescence spectroscopy.*

| **Analysis of Variance: 14.0 mg/L $TiO_2$ under 5.177 $\mu$W/$cm_2$/nm** | | | | | |
|---|---|---|---|---|---|
| **Source** | **DF** | **Sum of Squares** | **Mean Square** | **F Ratio** | **Prob > F** |
| DOC | 4 | 0.095084 | 0.023771 | 30.118800 | <.0001* |
| Error | 10 | 0.007892 | 0.000789 | | |
| C. Total | 14 | 0.102976 | | | |
| **Means for One-way ANOVA** | | | | | |
| **Level** | **Number** | **Mean** | **Std Error** | **Lower 95%** | **Upper 95%** |
| 0 | 3 | 1.1127 | 0.0162 | 1.0766 | 1.1489 |
| 1.57 | 3 | 1.0776 | 0.0162 | 1.0415 | 1.1138 |
| 2.95 | 3 | 1.0052 | 0.0162 | 0.9691 | 1.0414 |
| 4.28 | 3 | 0.9199 | 0.0162 | 0.8837 | 0.9560 |
| 5.71 | 3 | 0.9180 | 0.0162 | 0.8818 | 0.9541 |
| **Tukey-Kramer HSD for 14.0 mg/L $TiO_2$ under 5.177 $\mu$W/$cm_2$/nm** | | | | | |
| **q*** | **Alpha** | **Level** | | **Mean** | **Std Error** |
| 3.29108 | 0.05 | 0 | A | 1.1127 | 0.0162 |
| | | 1.57 | AB | 1.0776 | 0.0162 |
| | | 2.95 | B | 1.0052 | 0.0162 |
| | | 4.28 | C | 0.9199 | 0.0162 |
| | | 5.71 | C | 0.9180 | 0.0162 |

*Levels not connected by the same letter are significantly different
*Std Error used a pooled estimate for error variance
